# Biologically informed machine learning modeling of immune cells to reveal physiological and pathological aging process

**DOI:** 10.1101/2024.04.01.587649

**Authors:** Cangang Zhang, Tao Ren, Xiaofan Zhao, Yanhong Su, Qianhao Wang, Tianzhe Zhang, Boxiao He, Ling-Yun Wu, Lina Sun, Baojun Zhang, Zheng Xia

## Abstract

The immune system undergoes progressive functional remodeling from neonatal stages to old age. Therefore, understanding how aging shapes immune cell function is vital for precise treatment of patients at different life stages. Here, we constructed the first transcriptomic atlas of immune cells encompassing human lifespan, ranging from newborns to supercentenarians, and comprehensively examined gene expression signatures involving cell signaling, metabolism, differentiation, and functions in all cell types to investigate immune aging changes. By comparing immune cell composition among different age groups, HLA highly expressing NK cells and CD83 positive B cells were identified with high percentages exclusively in the teenager (Tg) group, whereas CD4_CTL precursors were exclusively enriched in the supercentenarian (Sc) group. Notably, we found that the biological age (BA) of pediatric COVID-19 patients with multisystem inflammatory syndrome accelerated aging according to their chronological age (CA). Besides, we proved that inflammatory shift-myeloid abundance and signature correlate with the progression of complications in Kawasaki disease (KD). Finally, based on those age-related immune cell compositions, we developed a novel BA prediction model, PHARE (https://xiazlab.org/phare/), which applies to both scRNA-seq and bulk RNA-seq data. Overall, our study revealed changes in immune cell proportions and function associated with aging, both in health and disease, and provided a novel tool for successfully capturing features that accelerate or delay aging.

## Introduction

Advanced age is often linked to increased morbidity and mortality from infectious diseases, and decreased vaccination efficacy^1, 2^. Aging is also a leading risk factor for the increased incidence of most cancer types^3, 4^. Although most children develop mild and self-limiting symptoms of infectious diseases, such as COVID-19, a severe and delayed post-SARS-CoV-2 inflammatory response in children has been recognized worldwide, correlating with disease severity^5, 6^. Moreover, compared with older individuals, children appear to have a more robust immune response during acute infections, whereas young cancer patients treated with immune checkpoint inhibitors often have poor outcomes^7,8^. The immune system exhibits highly age-specific features; however, the impact of age-related changes on different components of the immune system is not fully understood^2^. Therefore, the characterization of immune populations and function among different age groups could provide valuable insights into mechanisms underlying disease development and inform more precise disease treatments in the future.

Not everyone ages at the same rate. Usually, aging affects health and varies from person to person; therefore, it is not surprising that people with the same CA manifest diverse aging-related phenotypes^9^. In this regard, a person’s BA can often differ from his/her CA. Arrojo et al. revealed that most organs, even in one person, are also a mix of cells and proteins of vastly different ages, which depend on their rates of regeneration^10^. The ability to accurately measure human aging from molecular profiles has practical implications in many fields, particularly in disease prevention and treatment^11^. As expected, previous studies have developed many BA measurements that attempt to capture physiological changes during the aging process, such as DNA methylation, telomere length, and frailty^9, 11, 12^. Meanwhile, there are tools have been developed for BA prediction based on blood transcriptome, given the ease of obtaining samples^13, 14^. Considering that most immune diseases present strong age characteristics, the functional states of immune cells do not always match their CA in these diseases^15^. Therefore, effective measurements to comprehensively depict the age distribution of immune cells to accurately assess the impact of various diseases on the aging process are urgently needed.

Single-cell RNA sequencing (scRNA-seq) is a powerful technology used for studying individual cells and delineating complex cell populations. Recently, scRNA-seq has been performed in several studies to profile the immune landscape of human peripheral blood mononuclear cells (PBMCs) at different ages^16, 17, 18^. However, most of these studies have primarily focused on certain age periods, rather than the entire lifespan from 0 to over 110 years old, and many studies only analyzed a single type of immune cell, such as T cells^19, 20^, B cells^21^, or myeloid cells^22, 23^. Owing to the complexity of age spans and cell populations, the previous studies mentioned above limit our understanding of how immune profiles contribute to disease development from a systematic perspective. Thus, for the first time, we systematically analyzed all immune cells and their transcriptomic signatures in human PBMCs across different age groups encompassing the entire lifespan to establish an elaborate and aging-focused immune landscape.

In the current study, we first integrated three public scRNA-seq datasets from 24 healthy individuals across different age groups. Through extensive analysis, we revealed dynamic changes in cell composition, signaling pathways, metabolism, differentiation, and functions of human PBMCs over the lifespan. Furthermore, we re-analyzed other independent PBMC data from COVID-19 and KD mapping to healthy reference data, and detected a functional shift: immune cells’ BA differed from their CA, which was associated with the progress of vaccine efficacy and complications, respectively. Hereafter, we enrolled the largest PBMCs scRNA-seq datasets from 177 healthy individuals (over 1 million cells), and constructed a Physiological Age Prediction (PHARE) model. Collectively, our study provides essential insights for the precise treatment of patients at different life stages, and for building a novel machine-learning model to predict the patients’ BA at both scRNA-seq and bulk RNA-seq levels.

## Results

### Depicting global features of human immune cell atlas over the lifespan

To determine the effects of age on the immune system, we initially integrated three public scRNA-seq datasets of PBMCs from 24 healthy individuals across children (Cd), teenagers (Tg), adults (Ad), elders (Ed), and supercentenarians (Sc) (**Fig. 1a**). After quality control and filtering, we obtained high-quality single-cell transcriptomes from 159,671 cells (**Extended Data Fig. 1a and Supplementary Table 2**). Then, the scRNA-seq data were normalized and used harmony to remove batch effects (**Extended Data Fig. 1b**). Based on an unbiased integrative analysis across all immune cells, 12 cell clusters (**Extended Data Fig. 1c**) across six main immune cell populations were annotated according to the most salient cell markers: myeloid cells (*CD14^+^LYZ^+^VCAN^+^*), B cells (M*S4A1^+^CD79A^+^CD79B^+^*), T cells (*C3D^+^CD3E^+^CD3G^+^*), NK cells (*KLRD1^+^KLRB1^+^KLRF1^+^*), platelets (*PPBP^+^GP9^+^*), and hematopoietic progenitor cells (Hpc, *KIT^+^CD34^+^*) (**Fig. 1b, 1c, and Extended Data Fig. 1d**). Immune cell compositions were compared among different age groups, and the results showed that all six immune cell types were present in each group, albeit in different proportions. To confirm this validity, the proportion of cells in each individual was analyzed, and no immune cells showed high degrees of interindividual heterogeneity (**Fig. 1d**). As shown in **Figure 1e**, the composition of T cells significantly decreased, whereas that of NK and myeloid cells increased in young groups. Notably, the proportion of platelets gradually increased in old groups (**Extended Data Fig. 1e**), which is likely associated with an increased incidence of cardiovascular disease in these populations^24^. The compositions of immune cells from the elderly displayed prominent differences because the diversities measured with Shannon equitability index were significantly higher than those in young groups (**Fig. 1f**). The results implied different aging processes in different individuals, and this disparity was amplified with aging.

**Figure 1.**
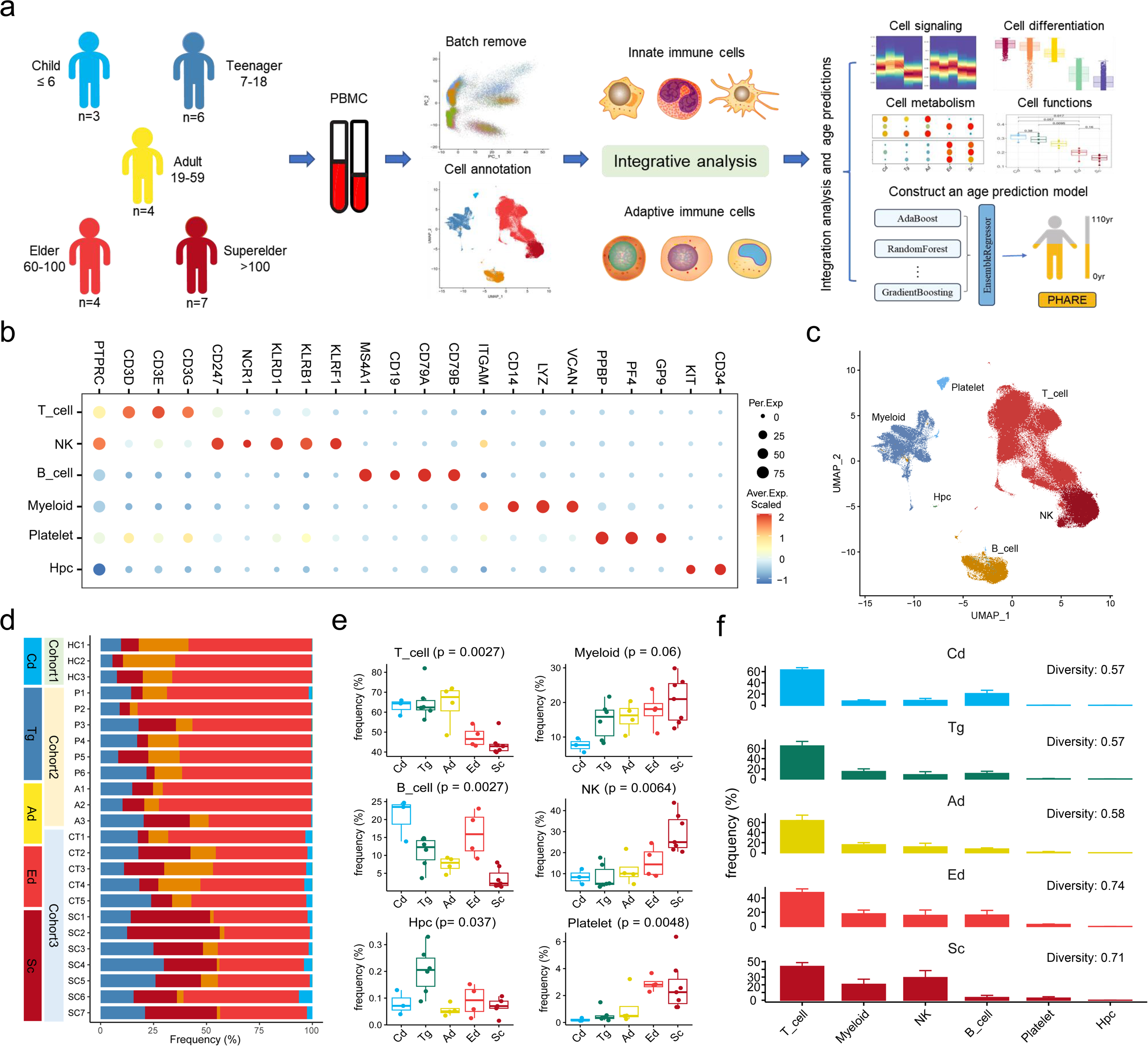
Depicting global features of human immune cell atlas over the lifespan. (**a**) Schematic diagram of the scRNA-Seq data collection, processing, and analysis design. (**b**) Dot plots showing representative signature gene expression in the main immune cell types. (**c**) UMAP projection of immune cell profiles in PBMC of different age groups. (**d**) Overview of data collection cohorts, clinical age groups; quantification of main cell types per patient and color-coded by main cell type. (**e**) Box plot showing distribution of main cell types across different age groups. The P values are calculated with kruskal.test. (**f**) Bar plots showing the main immune cell types from different age groups (mean ± SD). Average diversity measured with the Shannon equitability index for each tissue is shown. Point size of dot plot shows the fraction of cells with non-zero expression.

### Pluripotency functions in innate immune subsets decline with aging

Given that innate immune cells provide the first line of defense to control pathogenic infections and instruct the subsequent adaptive immune response, we analyzed the functional alterations in innate immune cells associated with aging. To classify each cell subpopulation in an unbiased manner, we re-clustered each cell lineage. Six cell clusters comprising five myeloid cell types with unique transcriptional features were revealed in the PBMCs of all age groups (**Extended Data Fig. 2a, and 2b**), including two types of CD14^+^ monocytes (*CD14_*Mono1 and *CD14_*Mono2), one group of CD16^+^ monocytes (*PCGR3A^+^*), and two types of dendritic cells (*CD1C^+^* cDC and *JCHAIN^+^* pDC) (**Fig. 2a and 2b**). Among the different age groups, we found that CD14_Mono1 was mainly enriched in the young groups (Cd, Tg and Ad), whereas CD14_Mono2 was markedly enriched in the elderly (**Fig. 2c and Extended Data Fig. 2c**). Indeed, in an independent PBMC bulk RNA-seq dataset, the CD14_Mono2 score of the elderly group (≥60) was significantly higher than that of the young group (<60), while the CD14_Mono1 score was significantly lower (**Fig. 2d)**. GSEA analysis of CD14_Mono1 and CD14_Mono2 DEGs found the former was enriched for migration- and response-related pathways, while the latter was enriched for epigenetic alterations and oxidative phosphorylation (**Fig. 2e and Supplementary Table 4)**. Consistent with previous findings^25^, both cDC and pDC decreased with increasing age (**Fig. 2c and Extended Data Fig. 2d**). We further evaluated the functional features of DCs in the different age groups. We found the genes involved in antigen presentation function of cDC gradually decreased with age, while the expression of splicing molecules increased (**Fig. 2f and Supplementary Table 5)**. With aging, the metabolic pattern of pDC was remodeled, while the maintenance of protein homeostasis was progressively lost (**Fig. 2f and Supplementary Table 5)**. To thoroughly explore the function of all myeloid subsets, we evaluated the module scores associated with well-defined signatures of inflammation regulation and immune activation (**Methods**). The results indicated that TNF and IL6 signaling were highly activated in *CD14_*Mono1 cells, while *CD16*_Mono cells showed high IFN-induced signaling (**Extended Data Fig. 2e**). For DC populations, pDC displayed strong protein secretion ability, whereas cDC had high MHC II expression and antigen processing potency (**Extended Data Fig. 2f**), which was in agreement with their respective roles. In addition, unlike CD14*^+^* monocytes, the proportion of CD16^+^ monocytes did not change substantially among different age groups (**Fig. 2c**). However, CD16*^+^* monocytes in the Tg group exhibited the highest levels of inflammation, IL6 and innate receptor signaling (**Extended Data Fig. 2g**). Taken together, the functions of monocytes and DCs gradually declined with aging.

**Figure 2.**
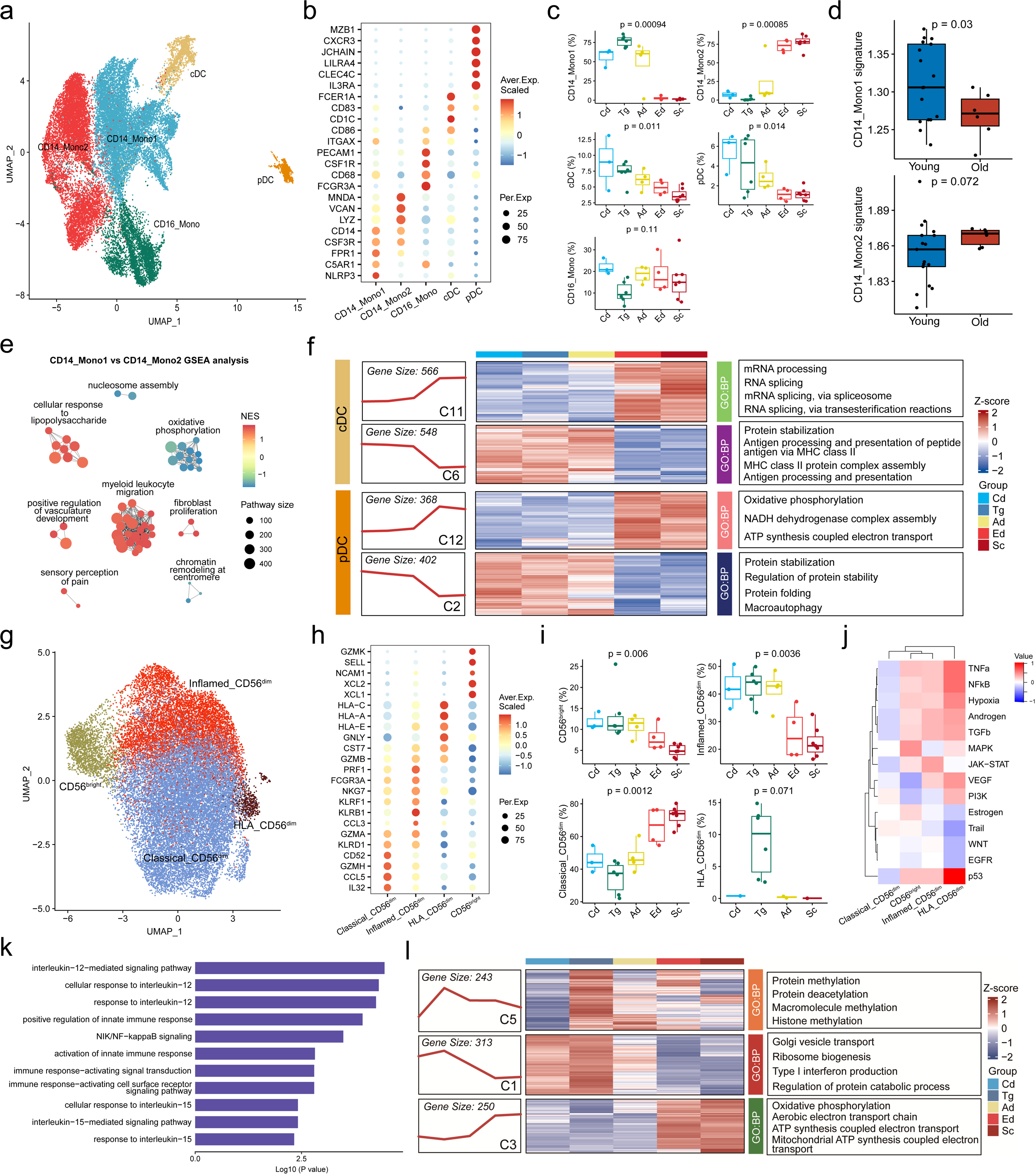
Pluripotency of functions in innate subsets declines with age. (**a**) UMAP projection of myeloid profiles in PBMC of different age groups. (**b**) Dot plots showing expression profiles of marker genes in different myeloid subsets. (**c**) Box plot showing distribution of myeloid subsets across different age groups. The P values are calculated with kruskal.test. (**d**) Box plot showing signature scores of monocyte subsets in different age groups based on ssGSEA algorithm, t-tests (two-sided) were performed. (**e**) Clustering network of significantly enriched GO pathways in the GSEA analysis. The nodes representing the significant GO pathways are colored by normalized enrichment score (NES). (**f**) Heatmap showing expression pattern of DC high-variable genes in different age group (left) and pathway enrichment analysis (right). (**g**) UMAP projection of NK profiles in PBMC of different age groups. (**h**) Dot plots showing expression profiles of marker genes in different NK subsets. (**i**) Box plot showing distribution of NK subsets across different age groups. The P values are calculated with kruskal.test. (**j**) Heatmap showing mean 14 PROGENy pathway scores of different NK subsets. (**k**) GO-based enrichment analysis illustrating indicated pathways upregulated in HLA_CD56^dim^ subset. (**l**) Heatmap showing expression pattern of NK high-variable genes in different age groups (left) and pathway enrichment analysis (right).

NK cells are innate immune cells that play critical roles in coordinating tumor immunosurveillance and viral infection^26^. Five clusters of NK cells were identified in the PBMCs of all the age groups (**Extended Data Fig. 3a**), and identified four NK cell types with unique transcriptional features (**Extended Data Fig. 3b**), including three types of CD56^dim^ NK (classical_CD56^dim^, inflamed_CD56^dim^, and HLA_CD56^dim^) and CD56^bright^ NK (**Fig. 2g and 2h**). We found that Classical_CD56^dim^ NK cells increased, whereas Inflamed_CD56^dim^ NK and CD56^bright^ NK cells decreased with aging (**Fig. 2i**). Notably, HLA_CD56^dim^ NK cells were specifically enriched in the Tg group (**Extended Data Fig. 3c**), although there was variation between healthy individuals (**Extended Data Fig. 3d**). Next, we used module scoring to evaluate the functional pathways related to NK based on the IOBR package, which provides a comprehensive investigation of the estimation of reported or user-built signatures^27^. Classical_CD56^dim^ NK cells had the lowest inflammatory signaling and CD56^bright^ NK cells had strong cytokine and chemokine secretion abilities (**Extended Data Fig. 3e**). Furthermore, we took advantage of PROGENy, a method that overcomes both limitations by leveraging a large compendium of publicly available perturbation experiments to evaluate 14 cell global functional pathways of NK subsets^28^. The result showed HLA_CD56^dim^ NK had strong inflammatory signatures and the p53 pathway was activated (**Fig. 2j**). Considering that HLA_CD56^dim^ NK specifically presented in the Tg group, we wanted to further explore its features. Pathway enrichment analysis of especially upregulated in HLA_CD56^dim^ NK cells revealed enrichment of terms associated with “response to IL-12”, “response to IL-15” and “NF-kB signaling”, which were consistent with the inflammatory phenotype (**Fig. 2k**). Consistent with the literature^29^, the CD56^bright^ NK population responsible for cytokine production also decreased with aging in our data (**Fig. 2i**). Additionally, the decrease in cytokine and chemokine secretion ability of this type of NK cell with aging was not due to cytokine regulation dysfunction but the decrease in golgi function and epigenetic alterations (**Fig. 2I**). Similar to CD56^bright^ NK cells, inflammatory CD56^dim^ NK cells also decreased with age, which may relate to cell cycle arrest (**Extended Data Fig. 3f**). Collectively, we revealed aging-related changes in NK cell composition and function.

### Age-group-specific adaptive subsets match distinct system immune function

Although the steps underlying the activation and differentiation of Age-engaged B cells have been extensively characterised^30, 31^, the cellular and molecular mechanisms underlying this process remain unclear. Therefore, we extracted B cells to further reveal seven cell clusters in PBMCs of all age groups (**Extended Data Fig. 4a**). Analysis of differentially expressed genes across these clusters revealed six B cell types with unique transcriptional features (**Fig. 3a and Extended Data Fig. 4b**): Naïve_B (*TCL1A^+^*), CD83_B (*CD83^+^*), Memory_B (*TNFRSF13B^+^*), Activated_memory_B (*CD86^+^*), Plasma_cell (*XBP1^+^MZB1^+^*), and Transitional_B cells (*CD5^+^*) (**Fig. 3b**). Unlike innate immune cells, B-cell subsets varied erratically across age groups except Plasma_cell (**Fig. 3c**). However, although the number of plasma cells gradually increased with age, the function of protein secretio declined (**Fig. 3d**). Notably, CD83_B cells were specifically enriched in the Tg group, and each patient in this group had a higher proportion of these type B cells (**Extended Data Fig. 4c and 4d**). To reveal the features of each B cell subset, we evaluated the module scores of functional pathways based on the signatures collected from previous studies. As depicted in **Extended Data Fig. 4e**, plasma cells showed higher protein secretion ability and lower antigen processing features among the six cell types, whereas CD83_B cells showed the highest inflammatory features. PROGENy analysis also illustrated that CD83_B cells had high TNF-α and NF-κB pathway scores (**Extended Data Fig. 4f**). Thereafter, we compared CD83_B cells with naïve B cells and found that Tg-enriched B cells upregulated activation-related molecules (*IGFBP4* and *CD69*) and inflammatory molecules (*CCL4* and *CCL4L2*) (**Fig. 3e and Supplementary Table 6**). Furthermore, gene set enrichment analysis (GSEA) similarly indicated that CD83_B cells were transcriptionally poised to IL-2 and IL-15 stimulated B cells in an inflammatory state (**Fig. 3f**). Aging is associated with decreased efficacy of vaccination in both humans and mice, with reduced B cell memory formation^32^. In this regard, we found that BCR signaling was significantly impaired in the elderly Memory_B cells, which was also related to the reduced efficacy of vaccination and increased susceptibility to infection in the elderly (**Extended Data Fig. 4g**). Activated memory B cells were enriched in the Ad group. To reveal this type of cell function, we compared activated memory B cells with memory B cells and found that the former upregulated many cytotoxic molecules, such as *NKG7, GNLY, CCL5*, and *GZMB* (**Extended Data Fig. 4h and Supplementary Table 7**). Taken together, our data suggest that B cells did not adhere strictly to age variation, and group-specific enriched subsets played special roles with distinct functional features.

**Figure 3.**
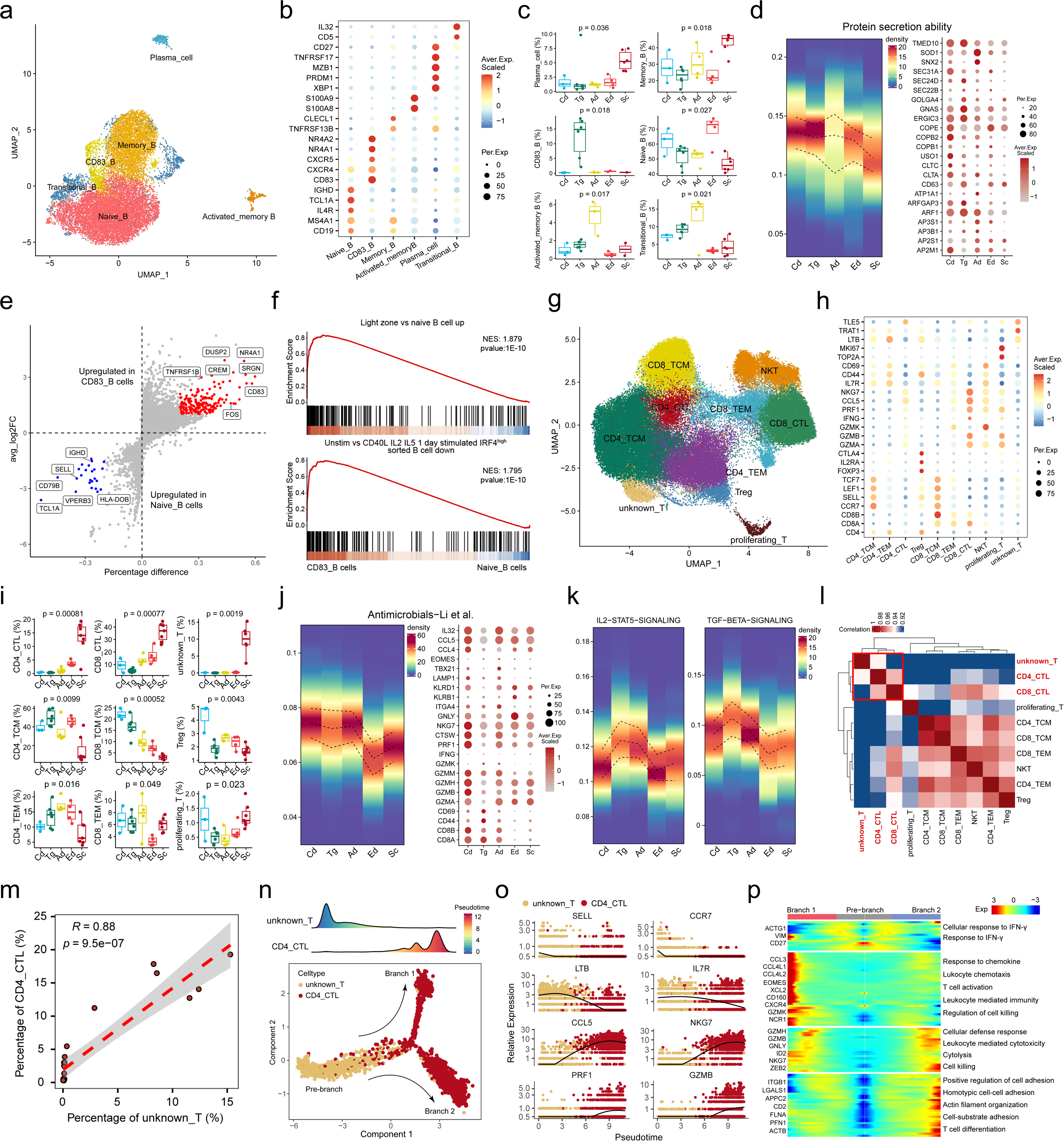
Age-group-specific adaptive subsets match distinct system immune function. (**a**) UMAP projection of B cell profiles in PBMC of different age groups. (**b**) Dot plots showing expression profiles of marker genes in different B cell subsets. (**c**) Box plot showing distribution of B cell subsets across different age groups. The P values are calculated with kruskal.test. (**d**) Densityheatmap showing indicates pathway score of plasma cells based on AUCELL algorithm (left), and representative genes dot plot (right). (**e**) Differential gene expression analysis using the log-fold change expression versus the difference in the percentage of cells expressing the gene comparing CD83_B cells versus Naïve_B cells (Δ Percentage Difference). (**f**) GSEA plots depict the enriched gene sets identified between the CD83_B cells and naïve_B cell subsets associated with B cell activation. (**g**) UMAP projection of T cell profiles in PBMC of different age groups. (**h**) Dot plots showing expression profiles of marker genes in different T cell subsets. (**i**) Box plot showing distribution of T cell subsets across different age groups. The P values are calculated with kruskal.test. (**j**) Densityheatmap showing indicates pathway score of CD8_CTL based on AUCELL algorithm (left), and representative genes dot plot (right). (**k**) Densityheatmap showing indicate pathway score of Treg based on AUCELL algorithm. (**l**) Heatmap showing transcriptomic similarity of T cell subsets. (**m**) Spearman correlation between percentage of CD4_CTL and unknown_T cells. (**n**) The distribution of T cell subtypes during the transition, along with the pseudo-time (upper). Subtypes are labeled by colors (lower). (**o**) Two-dimensional plots showing the dynamic expression of feature genes. (**p**) Heatmap showing the dynamic changes in gene expression along the different branches (left) and pathway enrichment results in each gene module (right). In figure **d**, **j** and **k**, the upper and lower curves of heatmap represent 75% and 25% of density, respectively. Point size of dot plot shows the fraction of cells with non-zero expression.

In contrast to B cells, there is a clear and well-accepted understanding that age-related aberrant T cell-driven cytokine and cytotoxic responses lead to the failure of immune tolerance and sensitivity to infectious diseases in older people^33^. Therefore, we extracted T cells to explore the timing and mechanisms behind the decline in T cell function. First, all T cells were clustered into 18 subsets (**Extended Data Fig. 5a**). We then identified four CD4^+^ T subsets (CD4_TCM, CD4_TEM, CD4_CTL, and Treg), three CD8^+^ T subsets (CD8_TCM, CD8_TEM, and CD8_CTL), NKT, proliferating_T, and unknown_T cells (**Fig. 3g**). Subsequent marker gene expression analysis confirmed the accuracy of these annotations (**Fig. 3h**), and the most up- and down-regulated genes were also calculated (**Extended Data Fig. 5b**). Among the T cell subsets, we found unprecedented percentage variation across age groups (**Fig. 3i**). Notably, CD4_CTL and unknown_T cells were especially enriched in the Sc group, although there was variation between patients (**Extended Data Fig. 5c and 5d**). To characterize various T cell subsets, we evaluated the functional pathways of all T cell subsets using module scores. Our results showed that, except CD4_CTL, CD4^+^ T cell subsets exhibited lower activities of HLA_signature and chemokine pathways than CD8^+^ T cell subsets (**Extended Data Fig. 5e**). Interestingly, CD4_CTL exhibited comparable cytotoxic and inflamed scores, yet they did not express the same exhausted features as CD8_CTL (**Extended Data Fig. 5f**). Furthermore, the PROGENy score also distinctively clustered CD4^+^ and CD8^+^ T cells (**Extended Data Fig. 5g**). CD8_CTL cells, which serve as the body’s robust defenders, were found to be most affected by aging^33^. We, therefore, examined the antimicrobial function of this T cell type across different age groups and found a decline in the elderly (**Fig. 3j**). Dot plots of related genes revealed that CD8_CTL cells in old people gradually lost the CD8 molecule (*CD8A* and *CD8B*) expression, whereas nature killer receptors (*KLRB1* and *KLRD1*) were upregulated. We also assessed the suppressive function of Tregs by analyzing the IL2-STAT5 and TGF-β pathways across different age groups. Results showed that although children had a higher percentage of Tregs, the Tregs in these periods were immature, possibly due to limited antigenic acclimation (**Fig. 3k**). Effector memory T cells (TEM), which indicate the body’s response to secondary immunity, were found to be immature in children, peaked in adults, and were diminished in the elderly, as determined by TCR signaling (**Extended Data Fig. 5h**). A similar pattern was observed in TCR and inflammatory signaling in NKT cells, while chemokine capacity was affected only by advanced age (**Extended Data Fig. 5i**). Recent research has highlighted the significant role of CD4_CTL cells in autoimmune diseases and tumor immunity, and an increase in these cells has been observed in supercentenarians^17, 34, 35^. However, the precursor cells for CD4 CTL remain unclear. As shown in **Figure 3I**, CD4_CTL and unknown_T cells were both enriched in the Sc group, prompting us to investigate a potential correlation between these two T cell types. Correlation analysis suggested that other T cells were most similar to CD4_CTL at the transcriptome level (**Fig. 3l**). Further correlation analysis revealed a significant strong correlation between the proportion of CD4_CTL and unknown_T cells (Pearson correlation value 0.88, **Fig. 3m**). According to previous study^36^, we discovered that unknown_T cells highly expressed molecules (*CD27, TCF7*, and *LTB*) of CD4 CTL precursors compared to CD4_CTL cells (**Extended Data Fig. 5j and Supplementary Table 8**). Therefore, we extracted CD4_CTL and unknown_T cells for pseudotime analysis to construct a developmental trajectory (**Fig. 3n**). We found that the expression levels of naive and stemness features were down-regulated, while killing molecules were gradually up-regulated along with pseudotime (**Fig. 3o**). We next investigated the transcriptional changes associated with two branches, and explored the functional features of two distinct CD4_CTL subsets. We identified one subset characterized by a response to chemokines, and another expressing adhesion molecules indicative of direct contact with target cell; both subsets acquired cell-killing abilities during differentiation (**Fig. 3p and Supplementary Table 9**). Together, we identified a type of CD4_CTL precursors (CD4_CTLP) that was enriched in Sc group, resulting in a higher percentage of CD4_CTL cells in the blood supercentenarians.

### Characterizing developmental hierarchies and quiescent immune cells using CytoTRACE

Considering that not all immune processes are uniformly sensitive to aging, various immune cell populations are heterogeneously influenced by the increase in age^37^. To reveal cellular states with intrinsic differences in function and developmental potential across different age groups, we used CytoTRACE to construct a cell differentiation atlas among PBMCs (**Methods**). As depicted in **Extended Data Fig. 6a**, all myeloid cells were divided into different cell clusters, which were highly consistent with the age groups according to their CytoTRACE scores. Additionally, the box plot showed that myeloid cells in the Cd, Ad, and Tg groups were less differentiated, while those in the elderly showed higher levels of differentiation. Surprisingly, myeloid cells in the Sc group exhibited a higher potential for differentiation capacity. The conclusion that aging markedly affects NK function remains controversial and is related to the diverse health statuses of the study subjects^38^. Our results demonstrated that NK cells in the Ad group had the highest differentiation capacity, suggesting that their differentiation function was well maintained into old age (**Extended Data Fig. 6b**). In contrast to innate immune cells, previous studies have indicated that adaptive immune function declined with age^30, 39^. However, our findings illustrated that the differentiation function of B/T cells did not decline in a straight line but gradually improved before the adulthood, peaked in adults, and then declined **(Extended Data Fig. 6c and 6d**). Unlike other immune cells, T cells in the Sc group were the most differentiated, in contrast to Ed, which strictly followed a trend of gradual decline with increasing age. Conversely, these results revealed that lymphoid and myeloid cells had different differentiation trajectories; the former had a well-established differentiation potential early on, while the latter developed this ability until adulthood.

### The single-cell metabolic profile reveals a shift in energy supply

Metabolism, being central to all biological processes, is crucial for the proper regulation of immune cells. Perturbations in metabolism can cause immune dysfunction and disease progression^40^. Therefore, we aimed to determine the metabolic activity of all immune cells across different age groups at single cell resolution. Following the instructions recommended by KEGG’s official website, we divided the extracted specific metabolic pathways into six categories (**Methods**). Lipid and carbohydrate metabolism declined the most dramatically with increasing age (**Extended Data Fig. 7a**). Notably, some lipid-related metabolic pathways showed an inverse trend, such as arachidonic acid metabolism in myeloid and NK cells, alpha-linolenic acid metabolism, steroid hormones, and fatty acid biosynthesis in B and T cells. As shown in **Extended Data Fig. 7b, oxidative phosphorylation activity was heightened in elderly groups across all immune cells, with other energy metabolism pathways also displaying increased activity in advanced age, excluding myeloid cells**. Amino acid metabolism is essential for metabolic rewiring, supporting various immune cell functions beyond increased ATP generation, nucleotide synthesis, and redox balance^41^. Notable amino acid metabolism-specific enhancements were observed in aged T cells. In particular, taurine and hypotaurine metabolism pathways were detected exclusively in T cells and were more active in the elderly. Intrigued by these findings, we further explored this metabolic pathway to differentiate between CD4^+^ and CD8^+^ T cell subsets. Surprisingly, it was not the CD8^+^ T cells (**Extended Data Fig. 7c)**, but the CD4^+^ T cells (**Extended Data Fig. 7d)** that showed an increasing trend with age. Collectively, these results highlighted that besides the uniformly active metabolites found in various immune cells of the elderly, there were also specific metabolites that were elevated exclusively in certain immune cells, which deserves further study.

### Decline of infection-fighting immune function with aging

To delineate the characteristics of entire immune cell subsets, we integrated all immune cells (Panage_data) and calculated the proportion of each cell type within all PBMCs across samples (Supplementary Table 3). Unsupervised hierarchical clustering, based on cellular composition, showed that the patients formed a remarkable divergence in different age groups (**Fig. 4a**). Odds ratio (OR) analysis revealed the cell distribution preferences of each age group, such as HLA_CD56^dim^ NK and CD83_B cells enriched in Tg and unknown_T cells, and CD4_CTL enriched in Sc (**Fig. 4b**). Furthermore, to confirm the preferences of HLA_CD56^dim^ and CD83_B cells in Teenagers and verify their widespread distribution, we incorporated another recently published scRNA-seq dataset from a study that included seven patients with multisystem inflammatory syndrome (MIS-C), eight patients with coronavirus disease 2019 (COVID-19), and seven age- and sex-matched healthy controls (HC)^42^. We mapped 124,448 immune cells from all patients to Panage_data, assigning cell annotation and age group labels based on transcriptome similarity (**Extended Data Fig. 6a and 6b)** (**Method**). From the OR analysis results, the HLA_CD56^dim^ and CD83_B cells also displayed a pronounced preference for the Tg group (**Fig. 4c**), while CD83_B and Treg cells were predominantly found in HC (**Fig. 4d**). These results suggested that CD83_B cells may confer protection against COVID-19, while reduced Treg content was associated with systemic inflammation. Considering the actual ages of all samples were below 18, we defined the immune cells mapped to Ad group as ‘shift-immune cells’. We then systemically assessed cell-type-specific functional changes among ‘shift-immune’ and ‘preserve-immune’ cells in COVID-19 and MIS-C cases (**Method**). We found that ribosome-related pathways were up-regulated in shift-immune cells, while TLR signaling, microbial and vaccine response-related pathways were significantly down-regulated (**Fig. 4e**). Besides, CD4_TCM cells underwent more significant remodeling in the MIS-C group than in the in COVID-19 group, while CD8_TCM cells were more profoundly affected in COVID-19 patients.

**Figure 4.**
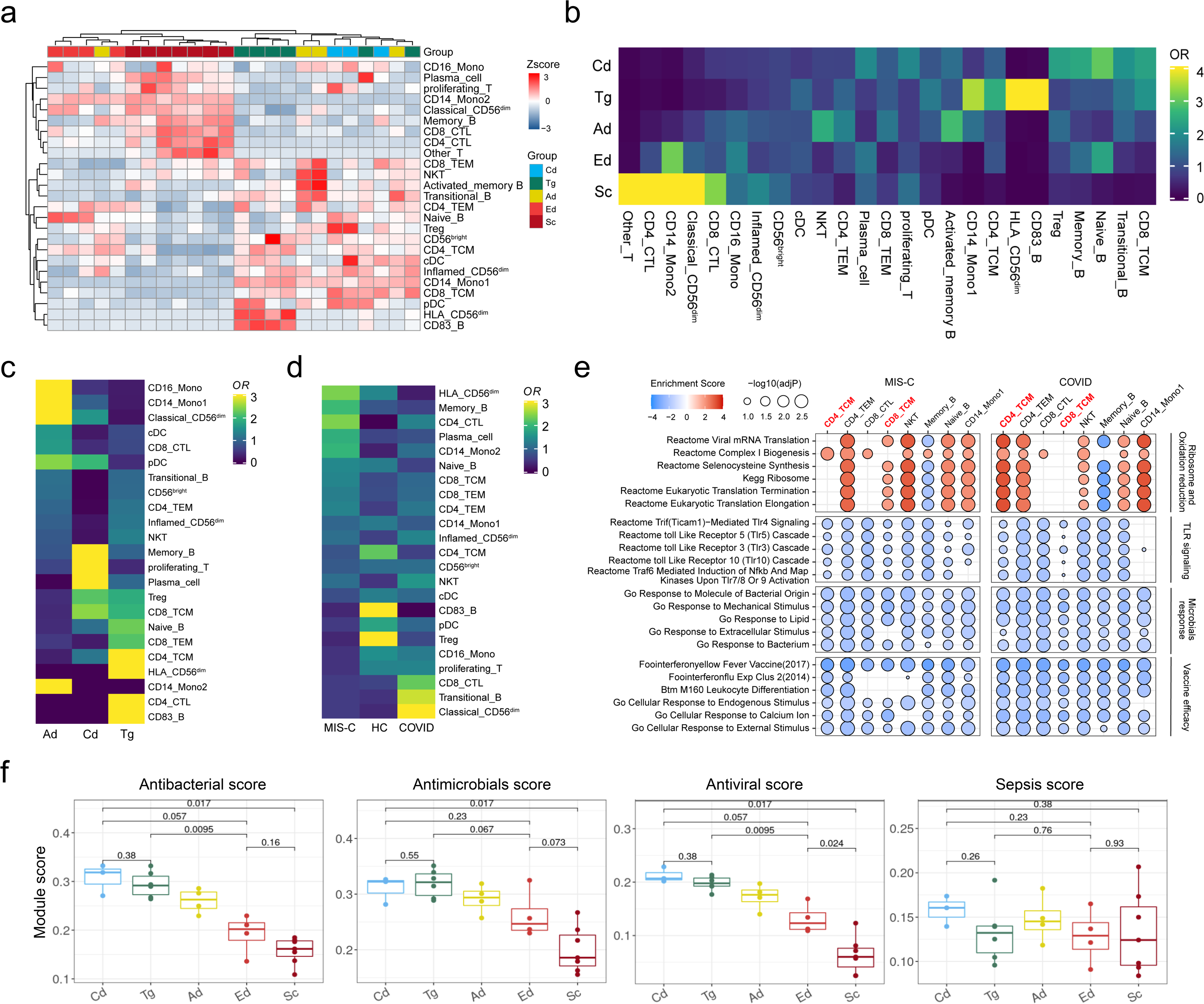
The anti-infectious functions gradually decline among age increase. (**a**) Heatmap showing clustering results based on the proportion of PBMC immune cell clusters, color-coded by age groups, patients indicated. (**b**) Heatmap showing the ORs of meta clusters occurring in different age groups based on TAIA data. Heatmap showing the ORs of predicted cell types occurring in different age groups (**c**) and clinical groups (**d**) based on independent COVID cohort. (**e**) GSEA of shifting versus preserving immune cells in MIS-C (left) and COVID-19 (right) patients. Selected gene sets are grouped into functional/pathway categories. Dot color denotes normalized gene set enrichment score, and size indicates log10(adjusted P value). (**f**) Comparing anti-infectious related module scores with different age groups based on each patient’s contribution. Statistical significance between groups is computed using a two-sided non-parametric Wilcoxon test.

Based on the differences in the composition of immune cells, we conducted a comprehensive assessment of the anti-infection ability and sepsis score for each age group based on all immune cells (**Methods**). As depicted in **Figure 4f**, except for the sepsis score, anti-bacterial, anti-viral, and anti-antimicrobial scores all declined progressively with increasing age. The cell types primarily contributing to these scores varied within each age group (**Extended Data Fig. 8c**). Considering the critical role of IFN signaling in anti-bacterial^43^, anti-viral^44^, and anti-antimicrobial^45^ abilities, we further evaluated two types of IFN signaling across different age groups. These signaling pathways consistently decreased in the two elderly groups (**Extended Data Fig. 8d**). In summary, we revealed that the immune cell composition of each age group was ever-changing, and that the capacity to combat infections depended on different immune cells.

### Functional shifting of inflammatory myeloid cells is associated with complication progression in Kawasaki disease (KD)

KD, a self-limiting vasculitis, predominantly affects children under 5 years old and has become the leading cause of acquired heart disease in children from developed countries^46^. Besides the primary febrile symptoms, KD can lead to serious arterial lesions, and the prompt diagnosis of KD remains challenging^47^. Moreover, a recent study has indicated that KD can accelerate immune cell senescence through the generation of reactive oxygen species^48^. Therefore, we examined whether scRNA-seq data from KD, mapped to Panage_data, could capture KD-induced cellular senescence. We reanalyzed scRNA-seq data from a previous study that collected seven patients with acute KD before and after IVIG therapy^47^. After data quality control, we obtained 35,336 high-quality cells for further analysis before IVIG therapy and mapping to Panage_data (**Extended Data Fig. 9a**). Considering that all KD patients were children (Cd), we defined the change from Cd to Cd (Cd-Cd) as the ‘preserve-state’ and all others (Cd-Tg, Cd-Ad, Cd-Ed, and Cd-Sc) as shift state according to the mapping result. We noted a remarkable divergence between the preserve- and shift-immune cell states, indicating transcriptomic difference between the two cell states **(Extended Data Fig. 9b**). Based on the aforementioned marker genes, six immune cell types were finely defined, including myeloid, NK cells, T/B cells, platelets, and Hpc (**Fig. 5a and Extended Data Fig. 9c**). Furthermore, we found varied shifting ratios across immune cell types, suggesting a broad influence of KD on immune cell senescence process **(Fig. 5b)**. To further verify the differential states of the shifting cells, all the immune cells were transferred to CytoTRACE. The result showed immune cells in the Cd group (preserve) possessed the highest degree of differentiation potential compared to the other groups (**Fig. 5c**). As depicted in **Figure 5d**, myeloid cells had higher shifting ratios (47.84%) than T (13.84%) and B cells (20.14%). Next, we assessed the functional differences between preserve- and shift-myeloid cells. GO enrichment analysis revealed that shifting myeloid cells increased the expression of genes involved in migration, cellular stress, and the inflammatory response (**Fig. 5e**). KEGG enrichment analysis identified pathways associated with severe KD complications, such as atherosclerosis, rheumatoid arthritis, and myocarditis in the highly expressed genes of shifting-myeloid cells (**Fig. 5f**). These findings suggest that abnormally elevated myeloid shifting in KD patients was the main cause of clinical symptoms. In addition, we also found that shifting-T/B cells all possess a more active state than preserve-cells (**Extended Data Fig. 9d and 9e**). Cell metabolism analyses showed that immune cells in the shifting state had higher lipid and vitamin-related metabolic activities in contrast to preserving cells, indicating that shifting cells presented an overwhelming immune response (**Fig. 5g and Extended Data Fig. 9f**).

**Figure 5.**
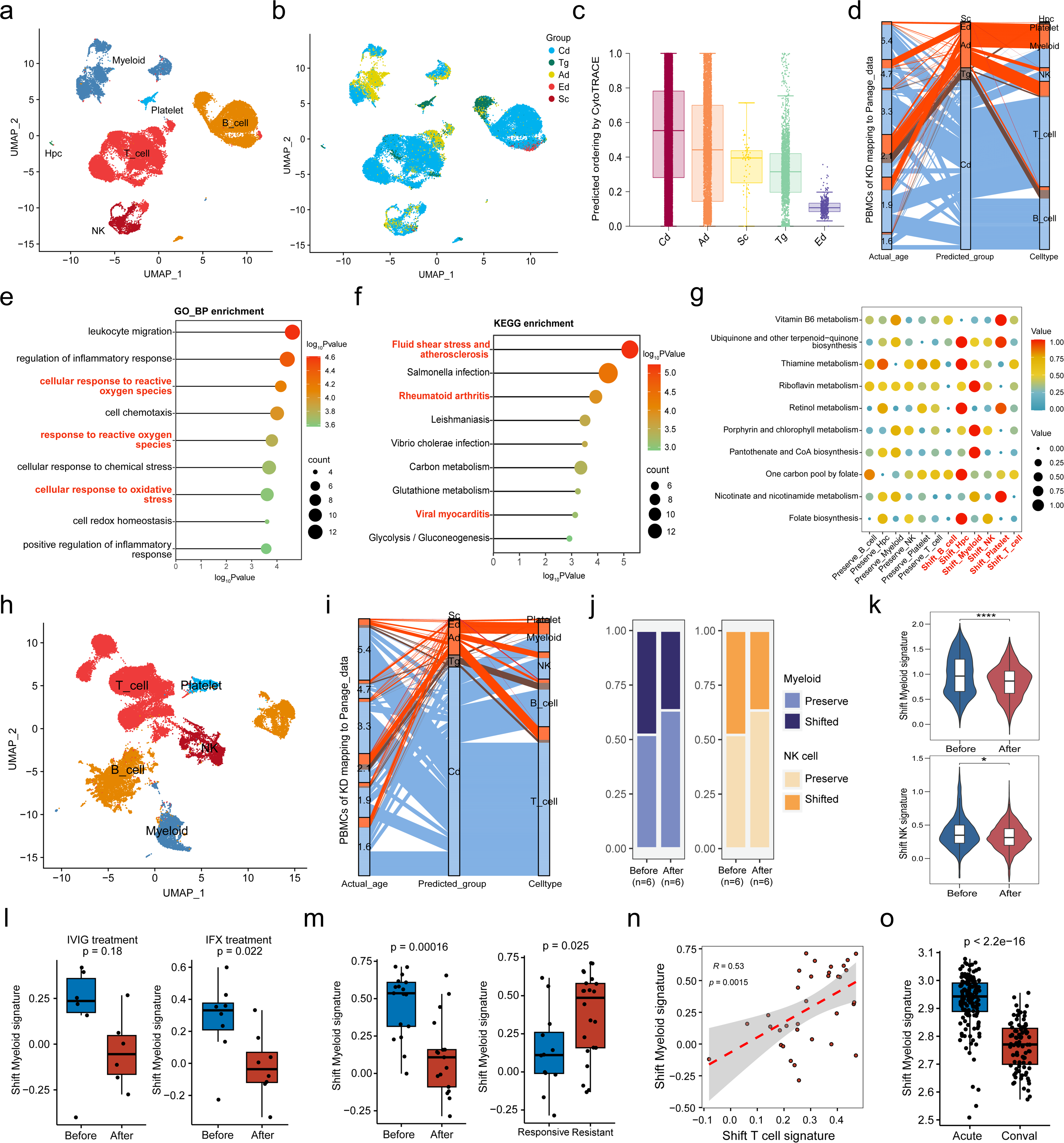
Functional-shifting inflammatory myeloid cells are associated with complications emerge in KD. UMAP projection of immune cell profiles from KD PBMCs, (**a**) group by cell types (**b**) and predicted groups. (**c**) Boxplots showing CytoTRACE values for different predicted groups of KD PBMCs. (**d**) Alluvial diagram showed the different shifting rates from KD PBMCs mapping to TAIA. Axis 1, 2, and 3 represented real age, predicted groups, and cell types. Dotchart showed GO-BP (**e**) and KEGG (**f**) based enrichment results of shift-versus preserve-myeloid upregulated genes. Color indicates log_10_Pvalue, and size indicates gene counts. (**g**) lipid-metabolic signature pathways in the main immune cell types of different age groups. Both color and size indicate the VISION score (**h**) UMAP projection of immune cell profiles from KD PBMCs after IVIG treatment, group by cell types. (**i**) Alluvial diagram showed the different shifting rates from KD PBMCs after IVIG treatment mapping to TAIA. Axis 1, 2, 3 represented real age, predicted groups and cell types. (**j**) Percentages of different myeloid and NK cell shift-subsets between before and after IVIG treatment, color-coded by cell subsets. (**k**) Violin plot showing shift-myeloid (upper) and NK (lower) signature score before and after IVIG treatment. The P values were calculated with wilcox.test. (**l**) Box plot showing shift-myeloid signature scores before and after IVIG treatment (left) or IFX treatment (right) based on ssGSEA algorithm. (**m**) Box plot showing shift-myeloid signature scores before and after IVIG treatment (left) or differential response groups (right) based on ssGSEA algorithm. (**n**) Spearman correlation analysis between shift-myeloid and T cell signature score. (**o**) Box plot showing shift-myeloid signature scores in different periods of onset based on ssGSEA algorithm. In figure l, m and o, the P values are calculated with t-tests (two-sided).

Intravenous immunoglobulin (IVIG) within the first 10 days after fever onset is the standard therapy for KD and remarkably reduces the rate of complications^49^. In this regard, we aimed to investigate the changes in KD PBMCs’ shifting ratios after IVIG therapy. We analyzed the scRNA-seq data of KD after IVIG therapy and mapped it to Panage_data (**Extended Data Fig. 9g**). The same immune cell types were identified based on marker genes (**Fig. 5h and** (**Extended Data Fig. 9h**). CytoTRACE analysis further confirmed that shift-cells exhibited a lower differentiation potential (**Extended Data Fig. 9i**). After IVIG therapy, each immune cell type was also found to shift-state, and the total shift-ratio was comparable (23.44% to 25.23%) (**Fig. 5i**). It is well established that monocyte abundance decreases and plasma cell numbers increase following IVIG therapy^47, 50, 51^. Our results further showed that the number of shift-B cells increased (20.14% to 36.10%), while shift-myeloid (47.84% to 36.62%) and NK cells (30.20% to 23.66%) decreased (**Fig. 5j**). Notably, shift-myeloid and NK signature remarkably down-regulated after IVIG therapy (**Fig. 5k**). Thereafter, we asked whether the shift-myeloid signature could be considered as a robust measure, and further evaluated changes after therapy using independent bulk RNA-seq datasets. We found shift-myeloid signature was significantly reduced after IVIG/IFX treatment (**Fig. 5i**). Besides, patients resistant to IVIG treatment had a more pronounced shift-myeloid signature compared to responsive patients (**Fig. 5m**). Correlation analysis revealed that shift-myeloid signature may also induce T cell senescence (**Fig. 5n**). It is known that KD progresses through different stages, but there is no diagnostic test for exact staging. We found that shift-myeloid signature was significantly higher in the acute phase than in the convalescent phase (**Fig. 5o**), which may provide new diagnostic insights for KD. Collectively, our results showed that the emergent pathogenic (shift) myeloid cells in PBMCs of KD patients were responsible for complications; therefore, treatments that attenuate its proportion and pathogenicity could alleviate symptoms.

### Constructing a machine learning predictor model to reveal the aging process

To build a transcriptome-based physiological age predictor model (PHARE), we compiled scRNA-seq datasets from PBMCs of healthy individuals spanning one lifespan, which contained more than one million cells (**Method**). All cells received cell annotation based on Panage_data, and then the proportion of all immune cell types was calculated to train the PHARE model (**Fig. 6a**). PHARE utilized two different pipelines for scRNA-seq and bulk RNA-seq data to predict physiological age from new datasets (**Fig. 6b**). We firstly evaluated the predictive performance of our model in the 177 healthy individuals using 10-fold cross-validation. Specifically, the individuals were divided into 10 roughly equal parts. In each iteration, the model was trained on 9 parts, with predictions made on the remaining part, and the mean Mean Absolute Deviation (MAD) between chronological age and predicted age was calculated as a performance metric. After completing 10 rounds of training and generating predictions for all individuals, the final average MAD all validations was 9.43 years (**Fig. 6c**). We noted there were no age differences were found when comparisons were made on each dataset, this result indicated the model was robust and independent of the different datasets (**Fig. 6d**). We then applied PHARE to new scRNA-seq datasets from various disease samples. As hypothesized, we found the predicted MAD for disease-derived datasets to be 13.61 years, higher than that for the healthy sample (**Fig. 6e**). Besides, we found MAD differed significantly across datasets from different disease sources, which indicated diseases had a variable impact on the aging process (**Fig. 6f**). Further analysis in systemic lupus erythematosus (SLE) patients revealed flare symptoms and those treated with no response had significantly higher MAD than those well-managed (**Fig. 6g**). This result suggested PHARE captured aging-accelerated features, which can help assess the impact of treatment. When applied to a COVID-19 dataset, PHARE revealed that patients with multisystem inflammatory syndrome in children (MIS-C) appeared older than ordinary COVID-19 patients and HC (**Fig. 6h**). This finding was consistent with the notion that hyper-inflammation can accelerate the aging process. Although PHARE was constructed based on scRNA-seq data, we wanted to further test whether it was also applicable for age prediction from bulk RNA-seq data. We then evaluated the accuracy of PHARE in two bulk RNA-seq datasets from healthy samples, which had a MAD between chronological age and predicted age of 9.09 years (**Fig. 6i**). This result indicated performance of PHARE on the bulk RNA-seq data remains robust. We further performed PHARE on another bulk RNA-seq data from coronary artery disease (CAD), and found CAD patients also had higher MAD than HC sample (**Fig. 6j**). Overall, these results illustrate that PHARE is a robust aging clock predictor, which can successfully capture aging-accelerated and aging-delayed features across various diseases.

**Figure 6.**
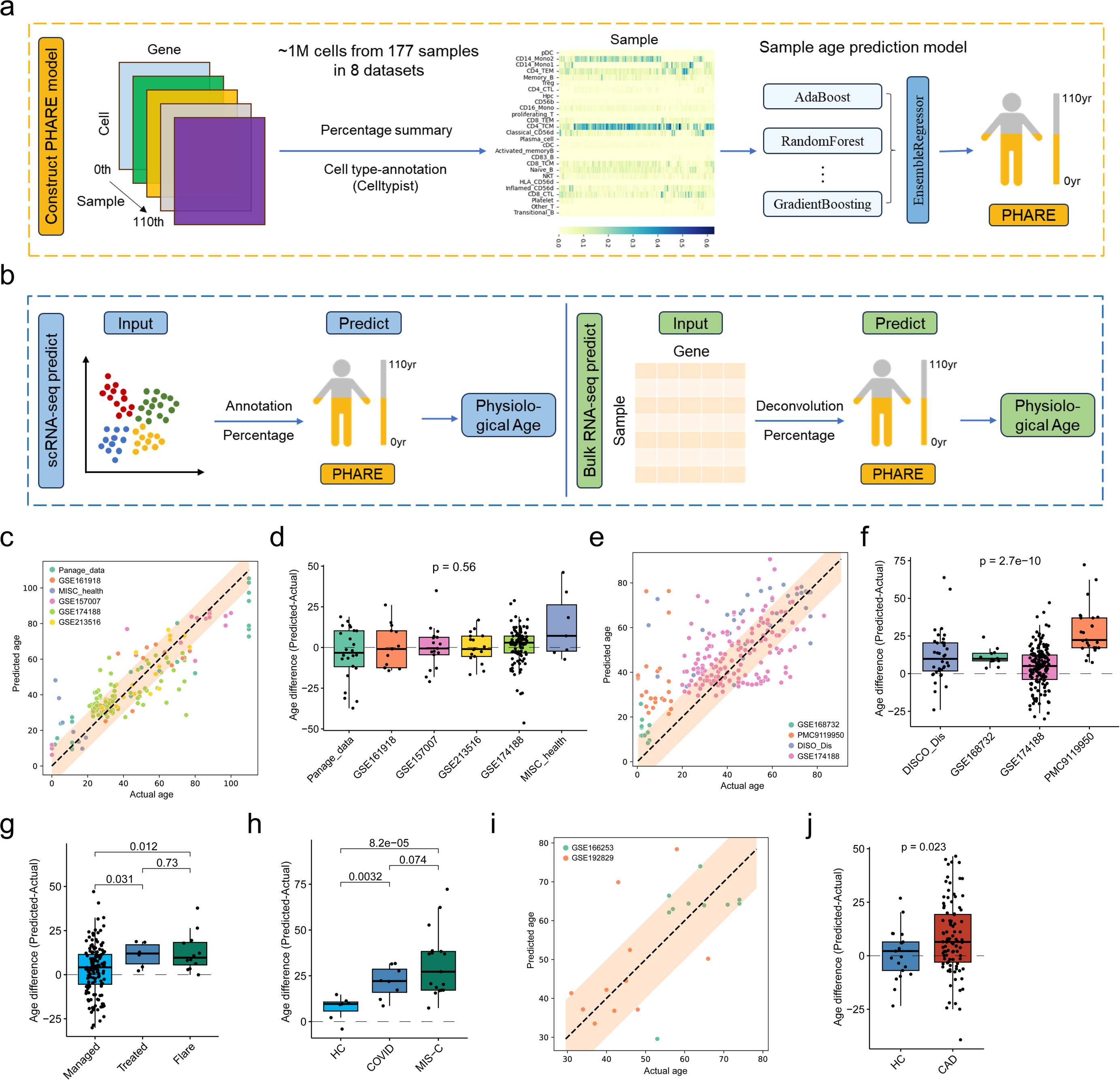
Constructing an age predictor model to explore human aging process. Workflow summary of PHARE construction (**a**) and applied PHARE to age prediction based on scRNA-seq and bulk RNA-seq data (**b**). (**c**) Spearman correlation analysis between predicted and actual age in scRNA-seq datasets from healthy people. (**d**) Box plot showing MAD was not different between different healthy scRNA-seq cohorts. (**e**) Spearman correlation analysis between predicted and actual age in scRNA-seq datasets from diseased people. (**f**) Box plot showing MAD was significantly different between different disease cohorts. (**g**) Box plot showing MAD was significantly different between different stages of SLE. (**h**) Box plots showing MAD were significantly different between different clinical groups. (**i**) Spearman correlation analysis between predicted and actual age in bulk RNA-seq datasets from healthy people. (**j**) Box plot showing MAD was significantly different between healthy and CAD samples based on bulk RNA-seq data, the P values are calculated with t-tests (two-sided). In figure c, d and f, the P values are calculated with kruskal.test. In figure j, h and j, the P values are calculated with t-tests (two-sided).

## Discussion

The aging process is complex, influenced by genetics, the environment, and their interplay. Given these complexities, traditional molecular biology approaches often fall short of elucidating the multimodal processes of aging. Single-cell techniques have emerged as powerful tools in high-throughput biology and systems immunology analyses^37^. By integrating single-cell datasets from previous studies^5, 17, 47^, our study illustrated immune cell signaling, metabolism, differentiation, and function changes of PBMCs, ranging from newborns to supercentenarians. We revealed several period-specific enriched cell types and reproduced these findings in independent datasets. Notably, we discovered a new population of CD4_CTL precursor cells in supercentenarians, the proportions of which are positively correlated with CD4_CTL, and differentiated into two distinct CD4_CTL subsets. By mapping immune cells from autoimmune and infectious diseases to healthy immune cells across human lifespan, we accurately captured aging-accelerated features at the cellular level. Collectively, our study constructed the first immune cell functional profile completely spanning people’s lifelong, and developed a machine learning model capable of predicting the BA of patients, suitable for both scRNA-seq and bulk RNA-seq datasets.

With aging, innate immune cells exhibit heterogeneous aging phenotypes, which correlate with their impaired capability for performing effective immune responses to newly encountered pathogens or vaccine antigens^29^. Consistent with a previous study, our results also showed a bias towards myeloid over lymphoid cell differentiation^52^. While the total percentage of myeloid cells was higher in the elderly, the proportions of both cDC and pDC decreased with age. Additionally, monocytes and dendritic cells progressively lose their immune stimulation function as they age. This contributes to why advanced age is associated with progressive immune deficiency, increased susceptibility to infections, and an impaired response to vaccination^53^. NK cells are another vital component of the innate immune system and serve as the first line of defense against infections and emerging malignancies. Our findings showed an increased percentage of NK cells in the elderly; however, a large proportion of these were senescent NK cells with weakened functional pathway signaling. Additionally, CD56^bright^ NK cell subsets, known for rapid proliferation and extensive cytokine and chemokine production upon activation, gradually decrease with aging^38^. Notably, we identified a new NK cell subset named HLA_NK cells, which were particularly enriched in the Tg group. Through pathway enrichment analysis, we found that HLA_NK cells showed high expression of IL-12 and IL-15 signaling genes. Previous studies have demonstrated that IL12 and IL15 sustain the effector function of NK cells in established tumors and infections^54, 55^. Therefore, this NK cell subset may contribute to the lower cancer rates and robust anti-infection abilities capabilities observed in teenagers.

Unlike the innate immune system, the functions of adaptive immune cells undergo drastic remodeling, often described as a shift from naïve to memory phenotypes with increasing age^2^. The representation of B cells is considerably altered with aging, and age-associated B cells, termed *CD19^+^CD21^-^CD23^-^* B cells in old mice and *CD19^+^CD38^−^CD24^−^* B cells in humans, were found to increase^56, 57^. In addition to the change in composition, B cell functions also become dramatically remodeled, gradually losing their antibody production capacity and acquiring inflammatory characteristics. We found a potential mechanism: a decreased protein secretion ability in plasma cells, marked by reduced expression of genes involved in protein trafficking (*ARF1* and *ARFGAP3*), protein sorting (AP-3 complex), and protein channel (*TMED10*) in the elderly^58, 59^. Unlike normal B cells, Age-Associated B Cells (ABCs) can rapidly respond to innate stimuli (toll-like receptors) and require minimal B-cell receptor (BCR) stimulation for activation^60^. Mechanistically, we found that BCR signaling in memory B cells, which are prevalent in the elderly, diminished with age. Additionally, CD83_B cells were specifically enriched in the Tg group, a result validated by an independent dataset. GSEA also indicated that CD83_B cells were transcriptionally predisposed to differentiate into light zone B cells. Considering that light zone B cells are activated and selected based on the affinity of their immunoglobulin, this finding implies that vaccination during adolescence might be more effective^61^. Owing to thymic involution in early adulthood, the maintenance of the naïve T-cell pool from the adult stage relies heavily on peripheral homeostatic proliferation^60, 62^. Moreover, because aging hematopoietic cells tend to favor myeloid differentiation, both our study and others have observed a gradual decline in total T cells^63^. Age most notably affects CD8^+^ T cells, characterized by a loss of naïve CD8^+^ T cells and gain of terminally differentiated CD8^+^ TEMRA cells (*CD45RA^+^KLRG1^+^CCR7^−^*)^64^. Contrary to a previous study, our data showed that CD8_CTL in the elderly did not highly express KLRG1 but rather KLRD1 and KLRB1, both of which acted as immune inhibitory receptors on NK cells. Peripheral homeostatic proliferation of T cells in humans effectively maintains the naïve CD4^+^ T cell pool, ensuring that CD4^+^ T cells are only moderately affected by age^60^. The mechanism by which CD4^+^ and CD8^+^ T cells are affected differently by aging remains unclear. Recently, CD4^+^ T cells with cytotoxic functions (CD4 CTL) have been recognized for their significant roles in viral infections, autoimmune diseases, and cancers^35, 65,66^. Hashimoto et al. illustrated that CD4 CTL dramatically increased in supercentenarians, a cell type also found in elderly patients with bladder cancer^17, 34^. Consequently, it is increasingly acknowledged that CD4 CTL have diverse functions and that their percentage is significantly associated with advanced age. On this basis, our results showed that CD4_CTL was detected in the peripheral blood of some adults, with a significant increase in the elders (60-110 yr), and the highest levels in supercentenarians (>100 yr). We further discovered that the emergence of CD4_CTLP (CD4 CTL precursors) in supercentenarians could be contributing to a sustained high generation of CD4_CTL in this group of people.

Biological aging is a complex process characterized by progressive deterioration occurring simultaneously at multiple levels. This process is tissue-specific and can even vary from one cell to another^9^. Accurate measurements to assess BA can be beneficial in capturing age-related physiological changes. Previous studies have developed numerous BA predictors through telomeres, epigenetic clocks, frailty, and other clinical biomarkers^67, 68, 69, 70^. However, all these measurements remained at the individual level, which significantly obscures the differences at single cell level. Therefore, we proposed a novel single-cell level BA prediction model (PHARE) based on the changes in every immune cell. Similar to other age prediction models^11, 13, 14, 71^, our approach also exhibits different MAD between the predicted ages and the patients’ actual ages. Upon analyzing KD (representing autoimmune diseases) and COVID-19 (representing infectious diseases) patient data, we consider the existence of MAD to be reasonable.

Collectively, our study has uncovered changes in immune cell function throughout the human lifespan, establishing a valuable reference for further research on aging. Our comprehensive analysis of the cell signaling, metabolism, differentiation, and functions of PBMCs provides an invaluable resource for the study of the aging process research, which will facilitate future precision treatments for diverse populations. Ultimately, we have developed PHARE into a user-friendly web tool (https://xiazlab.org/phare), which will greatly assist users without programming skills to use it for research on large-scale age-related diseases. In the future, we will continuously update PHARE and include additional samples from healthy individuals for training to enhance the accuracy of age prediction even further.

### Limitations of the Study

One limitation of our study was the use of a cross-sectional dataset in which associations can be identified rather than causalities. Further follow-up studies are required to validate these findings. In addition, sex-based differences in the immune system vary with age, and future studies need to include more samples to separate genders. Although we included data from multiple studies, we still did not define neutrophils and dendritic cells. A possible reason is that the 10x platform does not perform well on the capture rate of these two types of cells, and other single-cell sequencing platforms are required to explore these two types of cells in the future.

## Methods

### Panage_data collection and integration

To fully cover PBMC data across human lifespan, we included three separate studies. We extracted PBMC data from healthy children^47^ from Zhen Wang et al’ study, PBMC data from healthy teenagers and adults^5^ from Anjali Ramaswamy et al’ study, and PBMC data from healthy elders and supercentenarians^17^ from Kosuke Hashimoto et al’ study (Supplementary Table 1). Furthermore, we integrated cells into a shared space from different datasets for unsupervised clustering and used the harmony algorithm to correct the batch effect^72^. To detect the most variable genes used for harmony algorithm, we selected high-variable genes separately for each patient by “FindVariableFeatures” function from Seurat package (version 4.0.6). A consensus list of 2,000 high-variable genes was then formed by selecting the genes with the greatest recovery rates across patients, with ties broken by random sampling. Then, we ran Harmony with default parameters and integrated all datasets by using the “RunHarmony” function.

After dataset integration, the gene expression matrixes of all PBMCs were imported into Seurat v4 for subsequent analyses^73^. The following filtering steps were carried out to exclude low-quality cells: cells with fewer than 300 and more than 8000 detected genes were discarded; cells with a high fraction of mitochondrial genes (>10%) and hemoglobin genes (>1%) were removed. As a result, a total of 159,671 cells (Cd: 16,025, Tg: 57,945, Ad: 31,759, Ed: 15,716, Sc: 38,226) with a median of 1,087 genes were included in the further analyses. We applied SCTransform workflow (https://satijalab.org/seurat/) to analyze the scRNA-seq data with default parameters, which replaces the “NormalizeData”, “ScaleData” and “FindVariableFeatures” functions. Then, we performed a principal component analysis (PCA) dimensionality reduction (RunPCA) and selected the first 30 PCs to construct a shared nearest neighbor (SNN) graph (FindNeighbors). To visualize the clustering results, the non-linear dimensional reduction was performed with the Uniform Manifold Approximation and Projection (UMAP) method, and cluster biomarkers were found by the “FindAllMarkers” function from the Seurat package.

### External datasets mapping to Panage_data

To verify whether the cell types enriched in Tg group we identified in our origin datasets exist in other teenager PBMC samples, we included another published teenager PBMC scRNA dataset in our analyses^42^. For using Panage_data as a reference tool to interpret other datasets, PBMCs from KD patients^47^ and COVID-19 patients^42^ were included for further analysis. After the same data quality control, highly-quality immune cells were prepared for mapping to Panage_data. Then, marker genes common across a reference (Panage_data) and query (external datasets) were found by running FindAllMarkers, and the top 30 PCs were selected for cell clustering. Furthermore, we followed the tutorial (https://satijalab.org/seurat/v4.0/reference_mapping.html) to map each donor dataset from the query individually. We used the FindTransferAnchors function with reduction = “pcaproject” and MapQuery function as the previous study described^74^, and setting reduction = “pca” (as the documentation recommends for unimodal analysis). Finally, all PBMCs from external datasets were annotated according to our defined cell type.

### Scores quantification of function in scRNA-seq data

To score individual cells for Hallmark, KEGG, or other previously published functional pathway activities, we used multiple previously described methods to analyze different immune cell subsets. Firstly, the used human genes were from msigdbr (version 7.4.1) package, and gene sets were then used to score each cell. To eliminate the bias of sample background, we selected gene set enrichment analysis methods based on single-cell gene expression ranking AUCell^75^, UCell^76^, singscore^77^, and ssGSEA^78^. Of note, ssGSEA cancels the final standardization step, making it closer to the gene set enrichment analysis of a single cell. In addition, to evaluate whether the gene set is enriched in a certain cell subpopulation, we calculated the differential gene sets in the enrichment score matrix by Wilcox test (the filter criterion for differential genes is that the P-value after correction is less than 0.05). Finally, we used the rank aggregation algorithm (RRA) in the RobustRankAggreg package^79^ (version 1.1.0) to comprehensively evaluate the results of the different analyses, and screen out the genes that are significantly enriched in most gene set enrichment analysis methods Set (the filter criterion for comprehensive evaluation is P value less than 0.05).

### Anti-infection module score calculation

Module scores were calculated using the “AddModuleScore” function of Seurat with the default parameters. The anti-viral score consists of *SIGLEC1, RABGAP1L, IFI27, CADM1, RSAD2, MX1*, and *SERPING1*^80^. The anti-bacterial score consists of *SMPD1, CD44, SERPING1, SPI1, HERC1, MCTP1, FOLR3, CFAP45, PRF1, CTBP1, HLA-DRB1, ARL1, OAS3, ZER1, CHI3L1, IFIT2*, and *IFITM1*^80^. The anti-microbial score consists of 463 genes that come from Yongsheng Li et al’ study^81^. The sepsis score consists of *PLAC8, CLU, RETN, CD63, ALOX5AP, SEC61G, TXN,* and *MT1X*^82^. Besides, to reveal the anti-infectious mechanism, Type_I_IFN and Type_II_IFN related module scores were also calculated based on Rooney et al’ study^83^. In these four anti-infectious features, each patient has 25 cell types and each cell type has its specific score. For *i*-th patient, his/her patient’s score is calculated as follows:

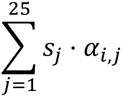

where *s*_*j*_ is *j*-th cell type’s specific score, and *⍺*_*i*,*j*_ denotes the proportion of the *j*-th cell of the *i*-th patient, satisfying:

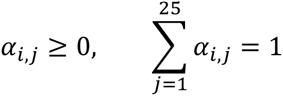

Then the differences between groups are compared for each patient of groups, the differential features in the established score matrix by Wilcox test. Other indicated functional gene sets used for cell annotation were extracted from IOBR (version 0.99.9) package^27^.

### Depicting cell global function using PROGENy

Considering lots of previous methods focused less on integrating responses from many different cell types, we chose PROGENy to address the accurate inference of pathway activity from gene expression in heterogeneous samples^28^. This method can overcome both limitations by leveraging a large compendium of publicly available perturbation experiments to yield a common core of Pathway RespOnsive GENes. Specifically, PROGENy evaluated 14 cell global functional pathways, which contained cell death, cell proliferation, cell metabolism, hormone signaling regulation and several immune signaling. Therefore, we used this method to evaluate the indicated immune cell function of different age groups.

### GO and KEGG Enrichment Analysis

Gene Ontology (GO) and Kyoto Encyclopedia of Genes and Genomes (KEGG) enrichment analyses were performed by ClusterProfiler (version 3.18.0) package, using genes specifically upregulated in indicated cell types as the input gene list. GO/KEGG biological process terms with Bonferroni-corrected P values (FDR) < 0.05 were considered significantly enriched terms. The aPEAR (version 1.0.0) package was used for clustering of enriched pathways and visualization. To cluster the differentially expressed genes, ClusterGVis (version 0.1.1) package was employed. Integration of the heatmap and functional enrichment analysis was also performed using the ClusterGVis package.

### Geneset and pathway scores assessment

To assign pathway activity estimates to individual cells, we applied GSVA using standard settings, as implemented in the GSVA package (version 1.42.0). For bulk RNA-seq data, we applied ssGSEA, as implemented in the IOBR package (version 1.0.0), to assess signature scores using the top 50 upregulated genes of indicated immune cell subsets (Supplementary Table 10). Gene Set Enrichment Analysis (GSEA) analysis was performed for each cell subpopulation using genesets (Hallmark) available at Molecular Signatures Database (MSigDB, https://www.gseamsigdb.org/gsea/downloads.jsp) or previously reported functional genesets^42^ with default parameters. P values were adjusted using the Benjamini–Hochberg method for the whole geneset list. Selected pathways shown in figure were manually curated to select genesets relevant to metabolism, infection and vaccine across the various differential expression comparisons.

### Detection of differentiation states based scRNA-seq data

Inferring both the state and direction of cell differentiation is challenging, scRNA-seq provides a powerful approach for reconstructing cellular differentiation trajectories. Notably, CytoTRACE, infers stem cell properties by scoring each cell based on its transcriptional diversity, which establishes a revelational RNA-based feature of developmental potential and a platform for the delineation of cellular hierarchies^84^. Therefore, we used CytoTRACE to reveal the differentiation degree of each cell type, which indicated the ability to respond to new antigens. The raw expression matrix (counts) for different immune cell subtypes was extracted and imported to the CytoTRACE package (version 0.3.3), and analyzed with default parameters. Then, the output CytoTRACE score for each cell was plotted with the “plotCytoTRACE” function, which visualized all cells in a low-dimensional embedding.

### Mining metabolic activity using scMetabolism

To estimate the metabolic activity of individual immune cells among different age groups, the scMetabolism (version 0.2.1) package was used developed by Yingcheng Wu et al^85^. We chose the KEGG database (https://www.genome.jp/kegg/pathway.html#metabolism) as the reference to quantify metabolic pathways in different cell types. In essence, selected KEGG metabolism pathways were generally divided into 6 categories (Amino acids, Lipids, Carbohydrates, Glycan, Cofactor/vitamin and Energy) according to the database recommendation. Considering VISION to be most appropriate for single-cell metabolism quantification among all methods, we evaluated the metabolism activity based on VISION score. Then, each metabolism activity was visualized using a modified “DotPlot.metabolism” function.

### Similarity and Correlation analysis

To measure the transcriptional similarity of T cell subsets, we evaluated the expression values of all genes in the T cell. Briefly, ward.D2 method was applied for hierarchical cluster analysis following Euclidean distance estimated by using the average expression of all genes. Then, spearman correlations were calculated between the different T cell subsets and visualization was performed by heatmap. For correlation analysis, the proportions of different immune-cell subsets or pathway scores were first calculated. Then, the correlation coefficients between them were estimated based on Spearman.

### Developmental trajectory inference of CD4_CTL cell

To characterize the developmental origins of CD4_CTL cells, we extracted unknown_T and CD4_CTL cells and imported them to the Monocle (version 2.14.0) software^86^ to construct developmental trajectory. The signature genes were calculated by dispersionTable function, and further filtered out low-expressed genes based on their expression values. Then, the CD4_CTL cell differentiation trajectory was inferred with the default parameters of Monocle after dimension reduction and cell ordering. After the cell trajectories were constructed, differentially expressed genes along the pseudotime were detected using the ‘‘differentialGeneTest’’ function.

### The odds ratio for cell distribution preferences

To characterize the cell distribution preferences, the odds ratio (OR) was calculated and used to indicate this feature as previously described^87^. OR is a measure of how strongly an event is associated with exposure and a ratio of two sets of odds. We first assumed that the cell type was *Ct* and the group was *g*, and calculated the OR to evaluate the preferences between *Ct* and *g*. The Fisher’s exact test was applied to this contingency table, thus OR and corresponding p-value could be obtained. Specifically, we defined a higher OR with a value > 2 indicating that indicated cell type *Ct* was more preferred to distribute in group *g*. P-values were adjusted using the BH method using the R function “p.adjust”.

### Diversity assessment based on immune cell composition

To compare the immune cell composition among different age groups, the frequency of immune cell meta-clusters was calculated separately. Then a Shannon equitability index (normalized Shannon diversity index) was calculated, and the details were described as previously reported^87^. The high index indicated composition of the various immune cells is more diverse.

### Physiological age prediction model construction (PHARE)

We developed a physiological age prediction model based on single-cell transcriptome data from normal peripheral blood samples (Supplementary Table 1). The specific steps are as follows: First, we used a single-cell dataset Panage_data that had been pre-annotated with various immune cell subtypes as a reference to train the Celltypist model^88^. The training process followed the standard procedure of Celltypist, which involves proportional sampling of each cell type from the reference dataset to enhance the ability to label rare cell types. Then, we used the trained model to label cell types across all other normal samples’ single-cell data. Ultimately, for each sample within all normal peripheral blood datasets (including the reference dataset), we calculated the proportion of each cell type and used these proportions as features to construct a sample age prediction model. This prediction model is an ensemble of three machine learning techniques: ’Random Forest’, ’Gradient Boosting’, and ’AdaBoost’, developed using the standard process of Sklearn^89^.

Subsequently, we developed a process to use the aforementioned trained model for physiological age prediction in samples at single-cell resolution and bulk resolution. For single-cell resolution transcriptome data samples, after cell type labeling using our trained Celltypist model, the ensemble model can directly provide age predictions (**Fig. 6b**). For bulk-resolution transcriptome data samples, we first employ cell type deconvolution technology, still using the pre-annotated single-cell dataset (Panage_data) as a reference, to parse out the cell type proportions in bulk samples. Specifically: We first use the *scanpy.tl.rank_genes_groups* function to calculate the up and down markers of the reference dataset; then, we retain markers unique to each cell type and use them with the intersection of genes contained in the bulk dataset as the reference genes for deconvolution; moreover, from the single-cell data, we calculate the average expression level of reference genes across each cell type to create a signature matrix; finally, we perform cell type deconvolution using the non-negative matrix factorization technique (NNLS), while providing NuSVR and Linear^90^ methods as alternative options for deconvolution algorithms. Through this process, we input the deconvoluted cell type proportions into the ensemble model, thereby providing age predictions for bulk resolution samples (**Fig. 6b**).

### Statistical analysis

Data were presented as means ± SEM with n per group and number of experimental replicates indicated in the respective figure legends. Statistical analyses were applied to biologically independent mice or technical replicates for each experiment which was independently repeated at least three times. Two-tailed Student’s t-test was used for statistical calculations based on bulk RNA-seq data, and wilcox.test was used for comparison based on scRNA-seq data. To assess differences between multiple groups, kruskal.test was performed. All bar graphs include means with error bars to show the distribution of the data. The level of significance is indicated as follows: *P < 0.05, **P < 0.01, ***P < 0.001, ****P < 0.0001.

## Supporting information

Supplementary Table 1. Detailed sample demographics of multiple cohorts utilized in this study

Supplementary Table 2. Absolute cell numbers for each cell type in each donor

Supplementary Table 3. The relative percentages for immune subsets in each sample

Supplementary Table 4. GSEA results of CD14_Mono1 vs CD14_Mono2

Supplementary Table 5. Enrichment results of gene patterns across different age groups in cDC and pDC

Supplementary Table 6. Enrichment results of gene patterns across different age groups in CD56-bright_NK

Supplementary Table 7. Differently Expressed Genes between CD83_B vs Naive_B cells

Supplementary Table 8. Differently Expressed Genes between other_T vs CD4_CTL cells

Supplementary Table 9. Enrichment results of different gene patterns in CD4_CTL differentiation along with pseudotime

Supplementary Table 10. Details of genesets generated in this study

## Code availability

The codes generated in this study are available at the GitHub repository (https://github.com/cliffren/PHARE).

## Data Availability Statement

All data needed to evaluate the conclusions in the paper are present in the paper and/or Supplementary Materials. Additional data related to this paper may be requested from the authors.

## Ethics Statement

The animal study was reviewed and approved by the Animal Care Committee of Xi’an Jiaotong University.

## Author Contributions

C.Z., Z.X., and B.Z. conceptualized the study and designed the article. C.Z., T.R., X.Z., and Y.S. analysed the scRNA-seq data. Q.W., T.Z., B.H., and L.-Y.W. provided insights into the analytic strategies. L.S., B.Z., and Z.X. supervised the study. C.Z., T.R., B.Z., and Z.X. wrote the manuscript with feedback from all other authors. All the authors read and approved the final manuscript.

## Fundings

This work was supported by grants from National Key Research and Development Program of China (2021YFA1100702), Major International (Regional) Joint Research Project (81820108017, B.Z.), National Natural Science Foundation of China grants (32170892 to B.Z.). National Natural Science Foundation of China (12231018 to L.-Y.W.).

## Acknowledgments

We thank Xiaoqi Wu from Genergy Biotechnology and Junan Yan from Shanghai BIOZERON for his help in technical support.

## Declaration of interests

The authors declare no competing interests.

## Supplementary figure legends

**Extended Data Fig.1.**
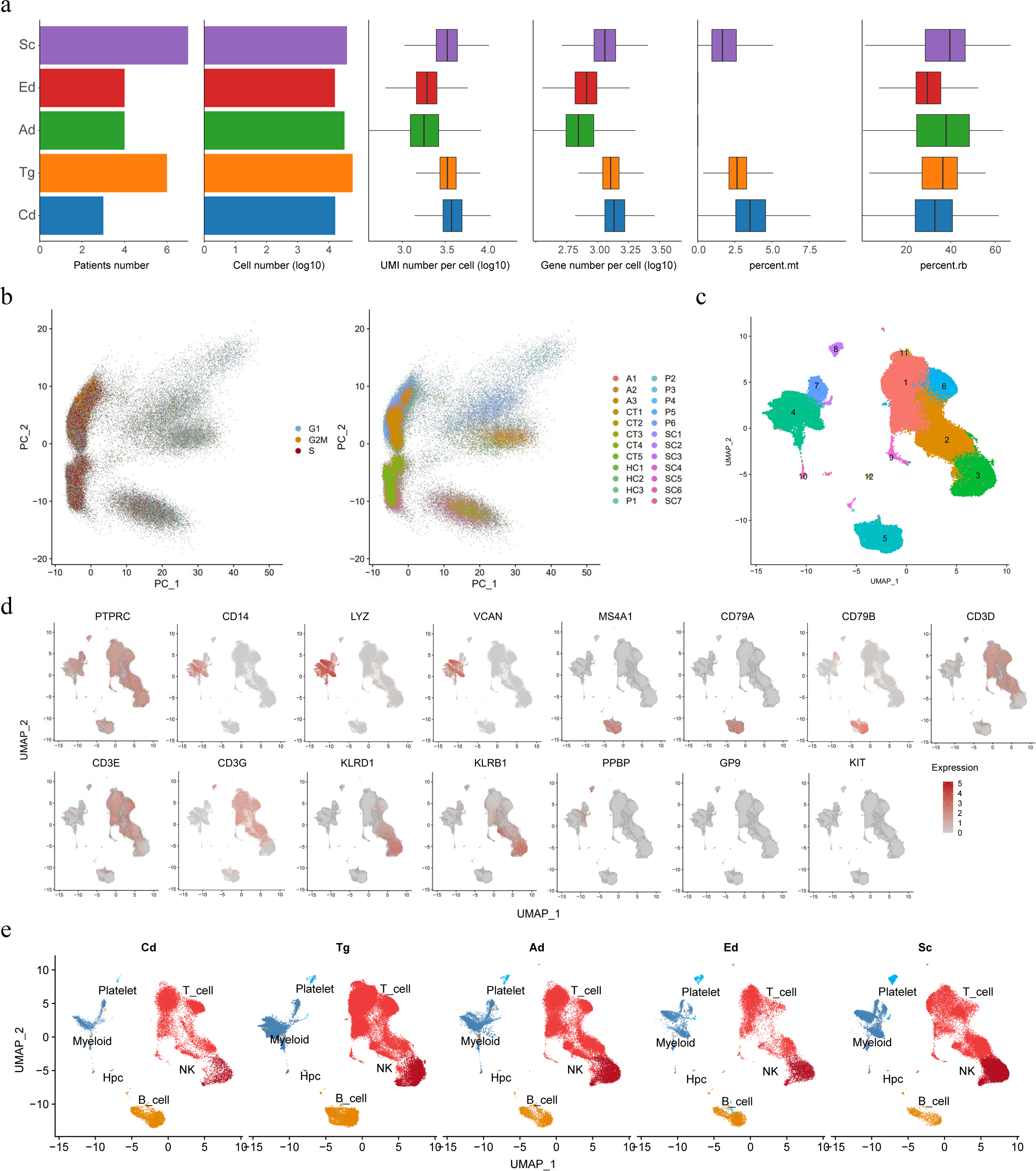
Quality control and cell types annotation. (**a**) Patient number, estimated cell number, mean reads per cell, gene number and ratio of mitochondrial/ ribosomal genes for scRNA-seq data across age groups. (**b**) PCA plot of 159,671 high-quality immune cells, colored by phase (left) or patients (right). (**c**) UMAP plot of the distribution of all immune cells, colored by cell clusters. (**d**) UMAP plots of scRNA-seq data representing the different immune cell clusters and selected marker gene expression. (**e**) All immune cells are projected onto UMAP visualization and split by age groups.

**Extended Data Fig.2.**
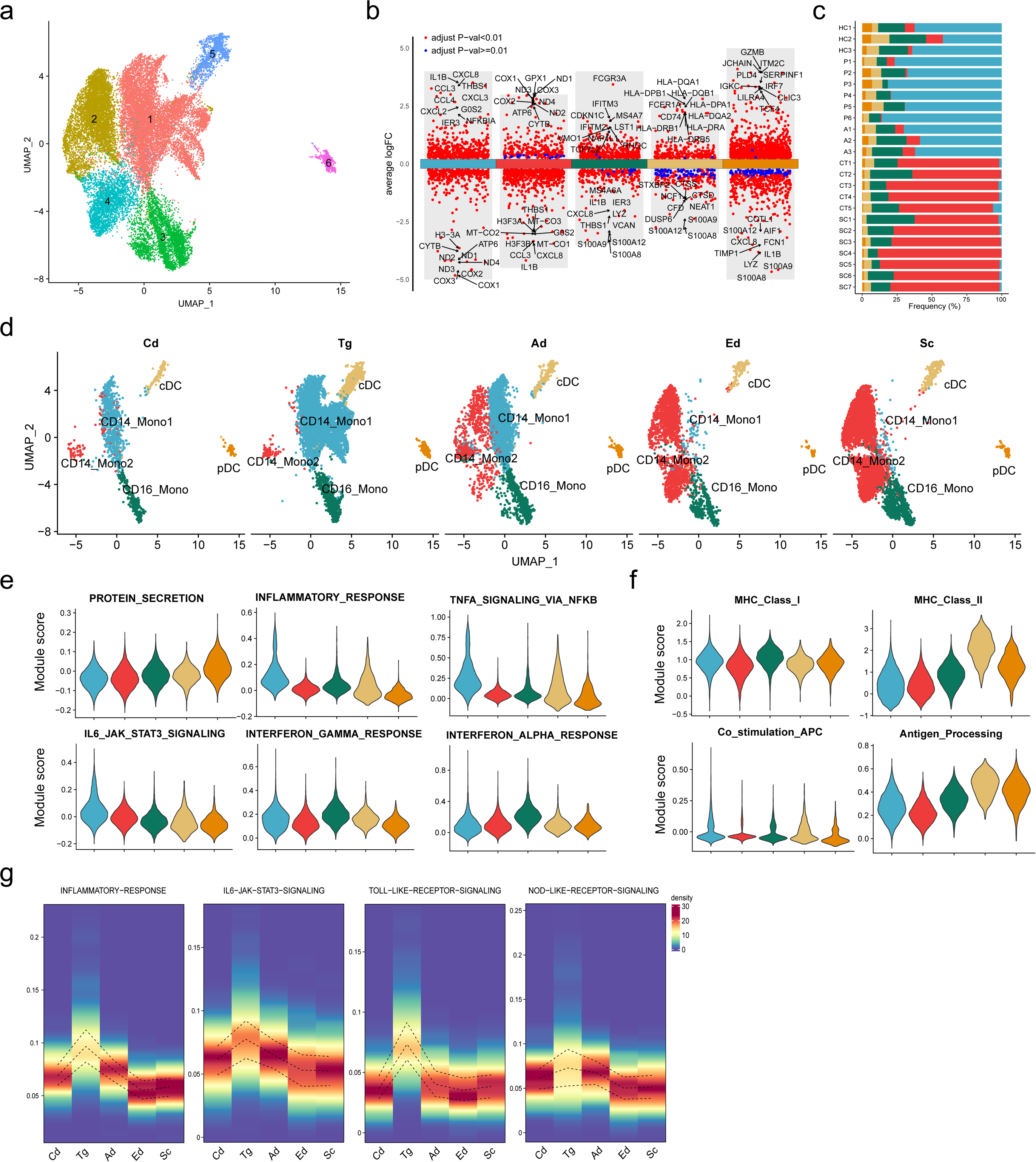
Single-cell transcriptomics revealed distinct myeloid subsets of different age groups. (**a**) UMAP plot of the distribution of myeloid cells, colored by cell clusters. (**b**) Differential gene expression analysis showing up- and down-regulated genes across all 5 myeloid subsets. An adjusted p value < 0.01 is indicated in red, while an adjusted p value > 0.01 is indicated in blue. (**c**) Quantification of myeloid cell types per patient, and color-coded by cell type. (**d**) Myeloid cells are projected onto UMAP visualization and split by age groups. Module scores of gene signatures related to (**e**) main features and (**f**) functional pathways of different myeloid cell subsets, for cell cluster color code see (**c**). (**g**) Densityheatmap showing indicates pathway score of CD14_Mono cells based on AUCELL algorithm of different age groups. The upper and lower curves of heatmap represent 75% and 25% of density, respectively.

**Extended Data Fig.3.**
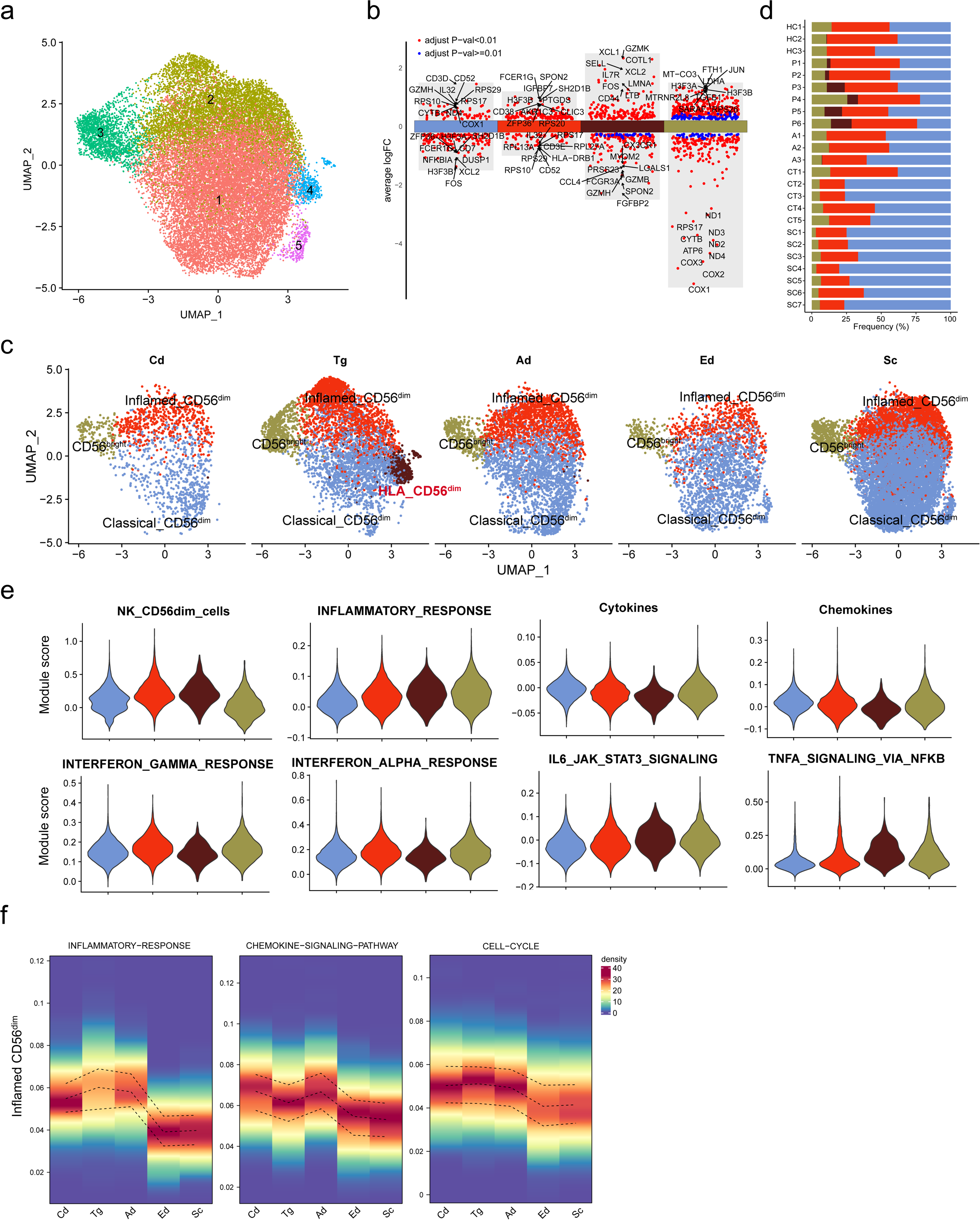
Single-cell transcriptomics revealed distinct NK subsets of different age groups. (**a**) UMAP plot of the distribution of NK, colored by cell clusters. (**b**) Differential gene expression analysis showing up- and down-regulated genes across all 4 NK subsets. An adjusted p value < 0.01 is indicated in red, while an adjusted p value > 0.01 is indicated in blue. (**c**) All NK projected onto UMAP visualization and split by age groups. (**d**) Quantification of different NK subsets per patient, and color-coded by cell type. (**e**) Module scores of gene signatures related to NK functional pathways of different myeloid cell subsets, for cell cluster color code see (**c**). (**f**) Densityheatmap showing indicates pathway score of Inflamed CD56^dim^ based on AUCELL algorithm. The upper and lower curves of heatmap represent 75% and 25% of density, respectively.

**Extended Data Fig.4.**
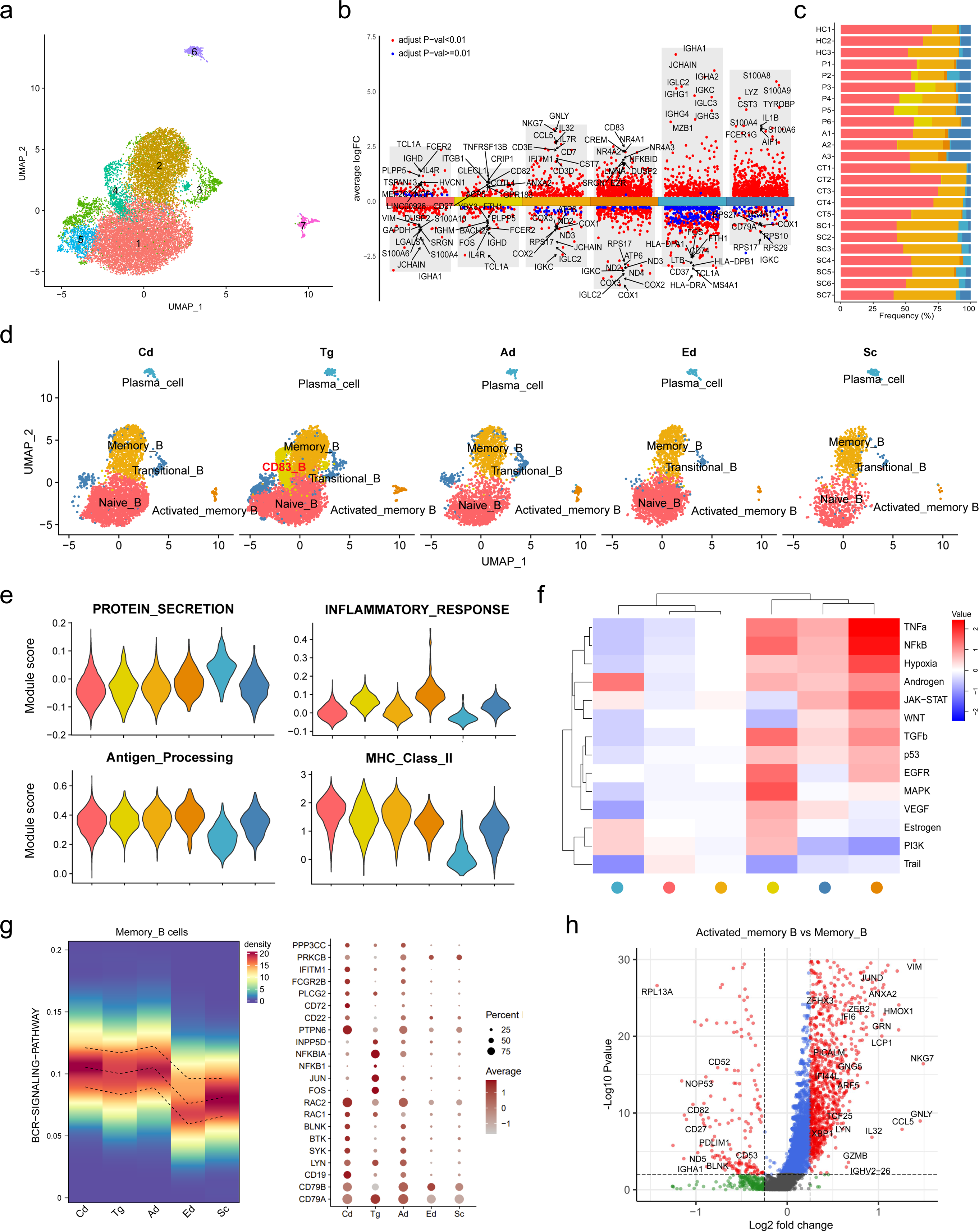
Single-cell transcriptomics revealed distinct B cell subsets of different age groups. (**a**) UMAP plot of the distribution of B cell, colored by cell clusters. (**b**) Differential gene expression analysis showing up- and down-regulated genes across all 6 B cell subsets. An adjusted p value < 0.01 is indicated in red, while an adjusted p value > 0.01 is indicated in blue. (**c**) Quantification of different B cell subsets per patient, and color-coded by cell type. (**d**) All B cells are projected onto UMAP visualization and split by age groups. (**e**) Module scores of gene signatures related to functional pathways of different B cell subsets, for cell cluster color code see (**d**). (f) Heatmap showing mean 14 PROGENy pathway scores of B cell subsets, for cell cluster color code see (**d**). (**g**) Densityheatmap showing BCR pathway scores of memory_B cells based on AUCELL algorithm (left), and representative genes dot plot (right). The upper and lower curves of heatmap represent 75% and 25% of density, respectively. (**h**) Volcano plots showing the DEGs of Activted_memory_B cells and memory_B cells subsets. The X-axis represents log2-transformed fold change. Y-axis represents −log10 transformed significance. Red points represent significantly expressed genes between these two groups.

**Extended Data Fig.5.**
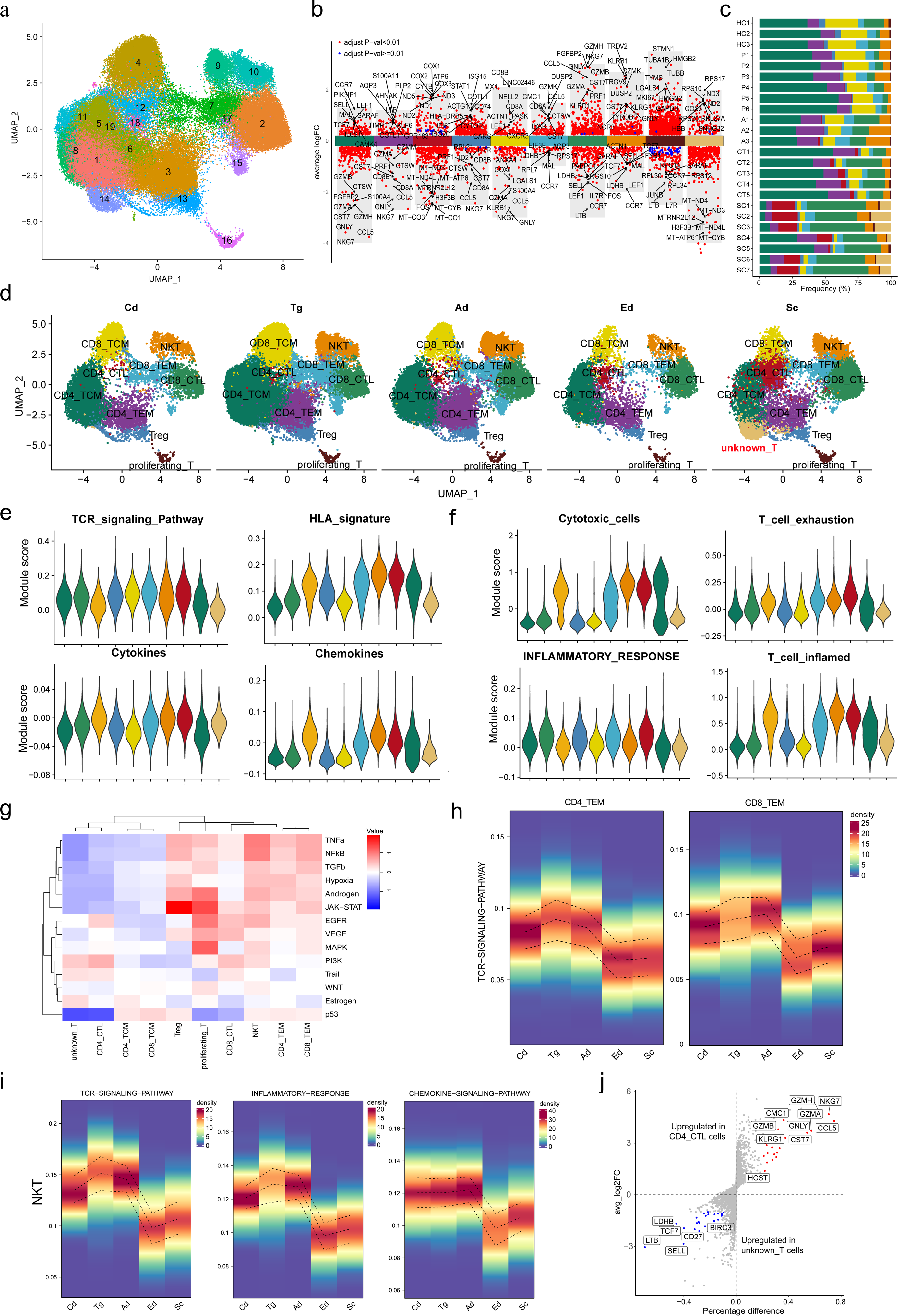
Single-cell transcriptomics revealed distinct T cell subsets of different age groups. (**a**) UMAP plot of the distribution of T cell, colored by cell clusters. (**b**) Differential gene expression analysis showing up- and down-regulated genes across all 10 T cell subsets. An adjusted p value < 0.01 is indicated in red, while an adjusted p value > 0.01 is indicated in blue. (**c**) Quantification of different T cell subsets per patient, and color-coded by cell type. (**d**) All T cells are projected onto UMAP visualization and split by age groups. Module scores of gene signatures related to (**e**) main features and (**f**) functional pathways of different T cell subsets, for cell cluster color code see (**d**). (**g**) Heatmap showing mean 14 PROGENy pathway scores across T cell subsets, for cell cluster color code see (**d**). (**h**) Densityheatmap showing TCR pathway scores of CD4_TEM and CD8_TEM cells based on AUCELL algorithm. (**i**) Densityheatmap showing TCR pathway and functional pathway scores of NKT cells based on AUCELL algorithm. (**j**) Differential gene expression analysis using the log-fold change expression versus the difference in the percentage of cells expressing the gene comparing CD4_CTL versus unknown_T cells (Δ Percentage Difference). In figure **h** and **j**, the upper and lower curves of heatmap represent 75% and 25% of density, respectively.

**Extended Data Fig.6.**
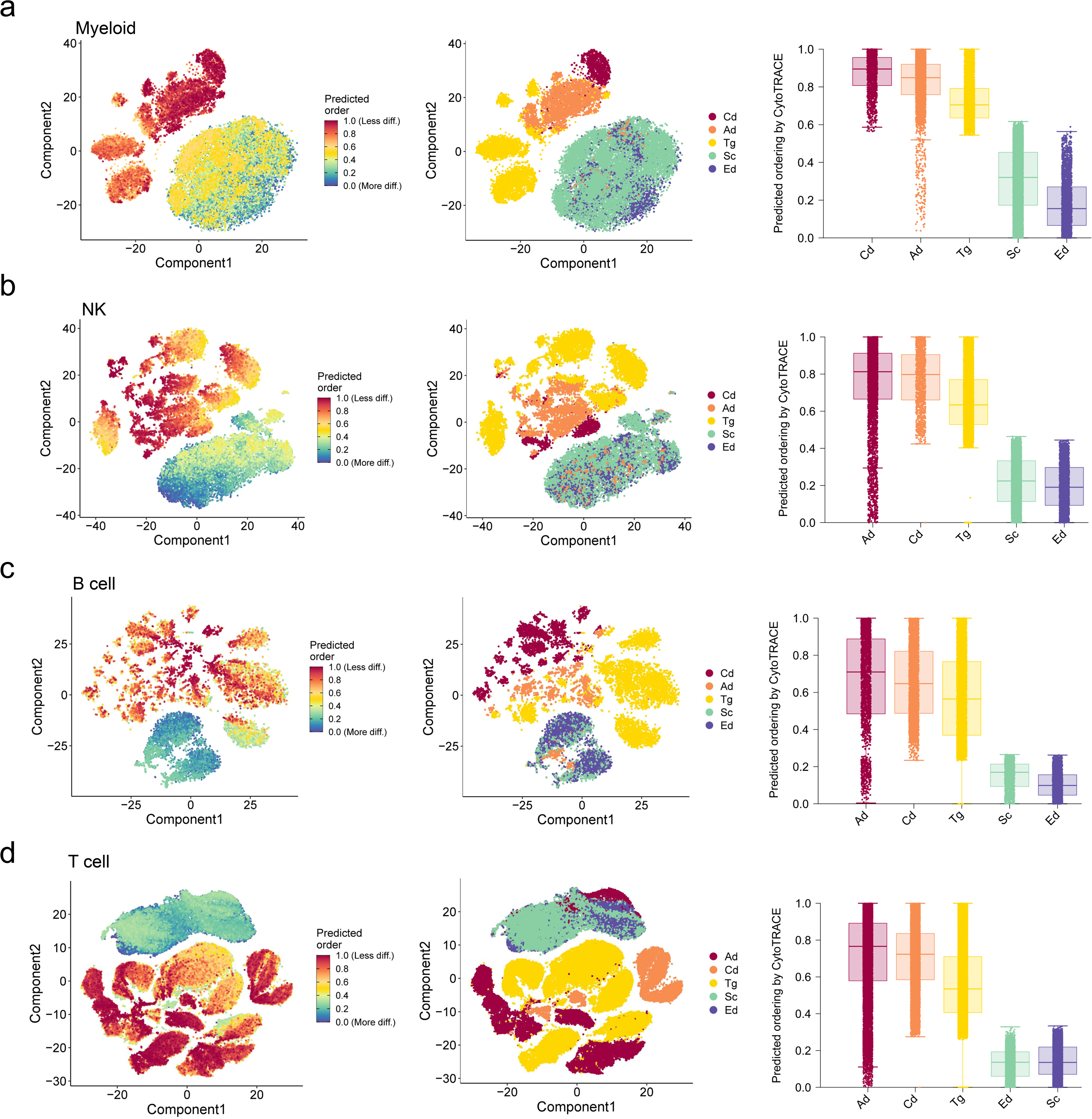
Characterization of developmental hierarchies and quiescent immune cells using CytoTRACE. (**a**) Low dimensional projections of myeloid cells were determined by CytoTRACE (left) and phenotype labels (middle) are depicted by color and velocity fields, and boxplots showing CytoTRACE values for candidate myeloid cells of different age groups (right). (**b**) Low dimensional projections of NKs were determined by CytoTRACE (left) and phenotype labels (middle) are depicted by color and velocity fields, and boxplots showing CytoTRACE values for candidate NKs of different age groups (right). (**c**) Low dimensional projections of B cells were determined by CytoTRACE (left) and phenotype labels (middle) are depicted by color and velocity fields, and boxplots showing CytoTRACE values for candidate B cells of different age groups (right). (**d**) Low dimensional projections of T cells were determined by CytoTRACE (left) and phenotype labels (middle) are depicted by color and velocity fields, and boxplots showing CytoTRACE values for candidate T cells of different age groups (right).

**Extended Data Fig.7.**
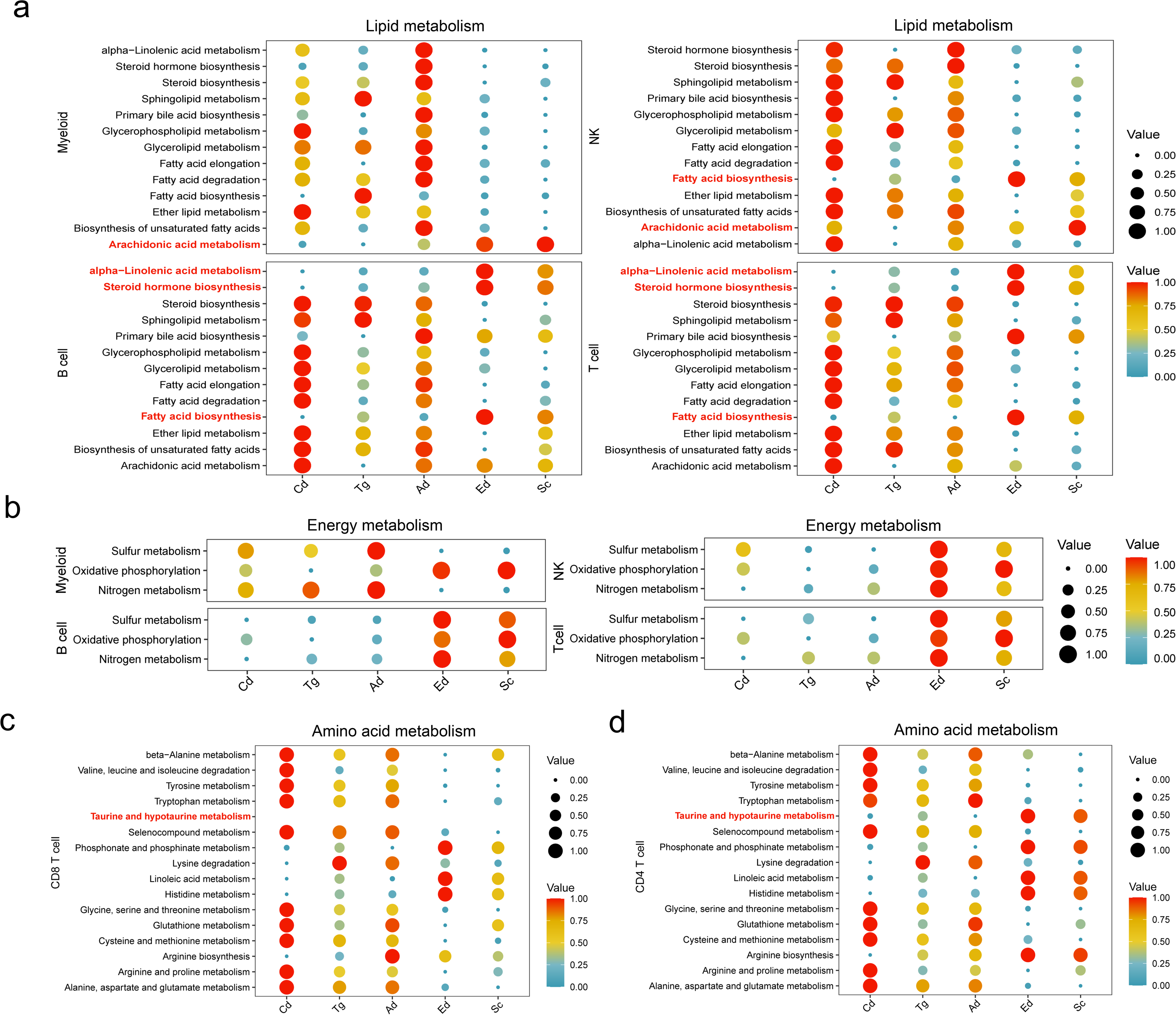
The single-cell metabolic profile reveals a shift in energy supply. (**a**) Dot plots showing intensity of representative lipid-metabolic signature pathways in the main immune cell types of different age groups. Both color and size indicate the VISION score. (**b**) Dot plots showing intensity of representative energy-metabolic signature pathways in the main immune cell types of different age groups. Both color and size indicate the VISION score. (**c**) Dot plots showing intensity of representative amino acids-metabolic signature pathways in the CD8^+^ T cells of different age groups. Both color and size indicate the VISION score. (**d**) Dot plots showing intensity of representative amino acids-metabolic signature pathways in the CD4^+^ T cells of different age groups. Both color and size indicate the VISION score.

**Extended Data Fig.8.**
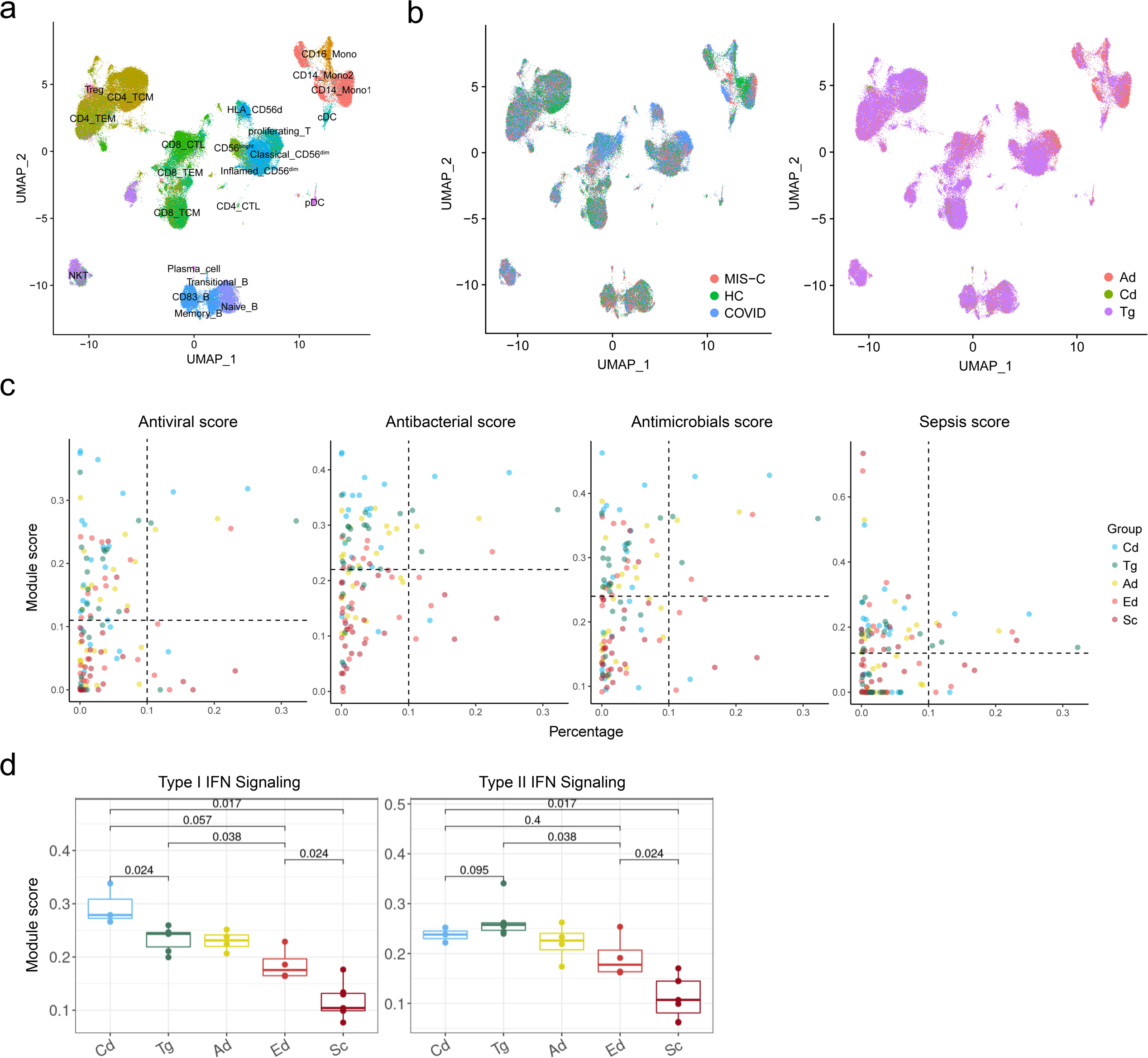
External validation of cell subsets enriched in teenager group. (**a**) UMAP plot of the distribution of immune cells from COVID-19 scRNA-seq dataset. (**b**) UMAP projection of predicted immune cell profiles from COVID-19 scRNA-seq dataset, colored by clinical groups (left) and predicted age groups (right). (**c**) Dot plots showing anti-infectious related module scores of different age groups, colored by age groups. Each point represents a cell subset. The X-axis represents percentage in all immune cell. Y-axis represents the indicated functional module score. (**d**) Comparing IFN-singling related module scores with different age groups based on each patient’s contribution. Statistical significance between groups is computed using a two-sided non-parametric Wilcoxon test.

**Extended Data Fig.9.**
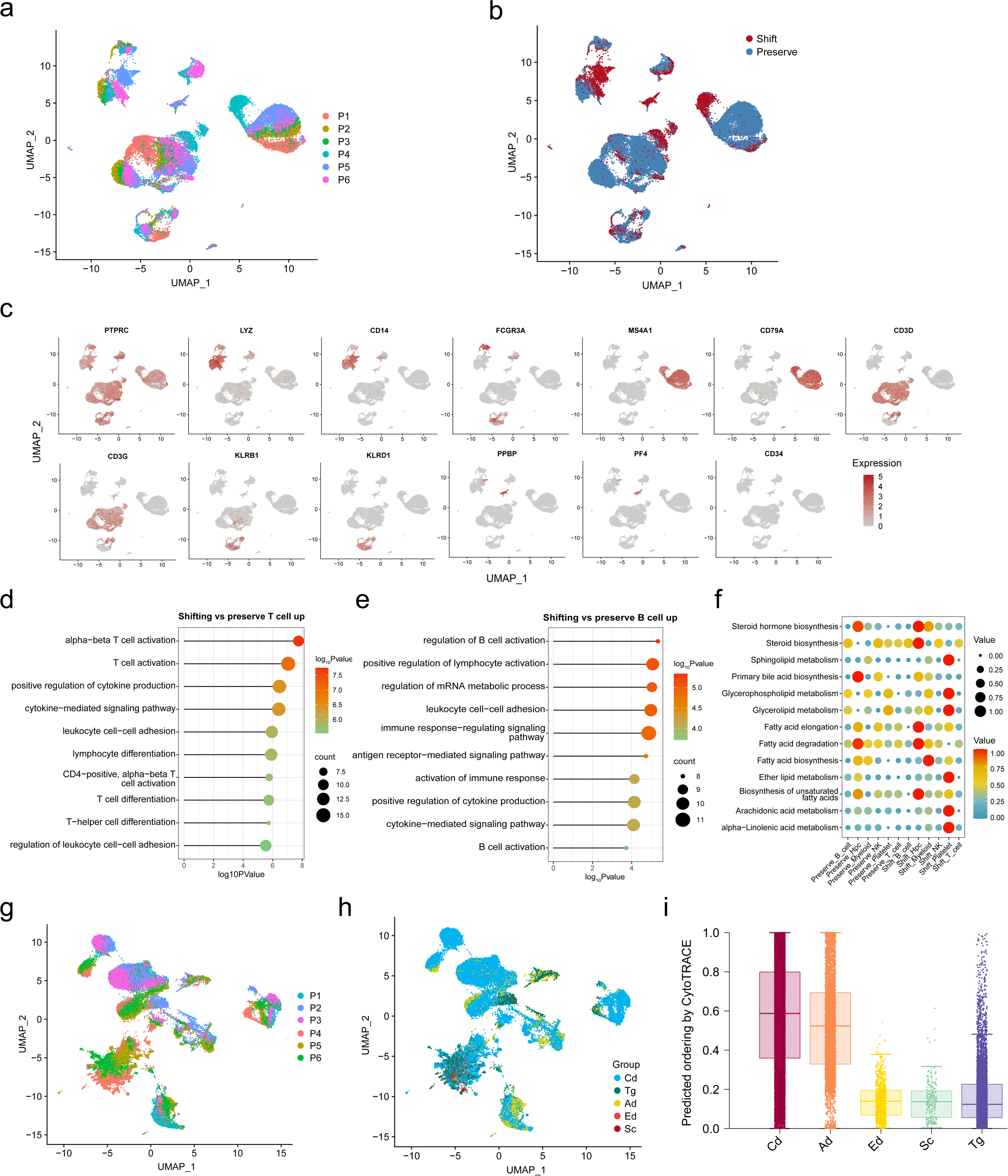
Reanalyzing previous KD PBMCs data based on TAIA as reference. (**a**) UMAP projection of immune cell profiles from KD PBMCs, group by patients (left) and predicted states (right). (**b**) UMAP plots of KD PBMCs data representing the different immune cell clusters and selected marker gene expression. (**c**) UMAP plots of scRNA-seq data representing the different immune cell clusters and selected marker gene expression. (**d**) Dotchart showed GO-based enrichment results of shift-versus preserve-T cells upregulated genes. Color indicates log_10_Pvalue, and size indicates gene counts. (**e**) Dotchart showed GO-based enrichment results of shift-versus preserve-B cells upregulated genes. Color indicates log_10_Pvalue, and size indicates gene counts. Dot plots showing intensity of representative. (**f**) lipid-metabolic signature pathways in the main immune cell types of different age groups. Both color and size indicate the VISION score. UMAP projection of immune cell profiles from KD PBMCs after IVIG treatment, group by (**g**) patients and (**h**) predicted groups. (**i**) Boxplots showing CytoTRACE values for different predicted groups of KD PBMCs after IVIG treatment.

## Notes

https://xiazlab.org/phare/

### Competing Interest Statement

The authors have declared no competing interest.

### Summary of Updates

We found some errors in the author order and spelling, and the new version has corrected them.

